# Object-vector coding in the medial entorhinal cortex

**DOI:** 10.1101/286286

**Authors:** Øyvind Arne Høydal, Emilie Ranheim Skytøen, May-Britt Moser, Edvard I. Moser

## Abstract

Mammals use distances and directions from local objects to calculate trajectories during navigation but how such vectorial operations are implemented in neural representations of space has not been determined. Here we show in freely moving mice that a population of neurons in the medial entorhinal cortex (MEC) responds specifically when the animal is at a given distance and direction from a spatially confined object. These ‘object-vector cells’ are tuned similarly to a spectrum of discrete objects, irrespective of their location in the test arena. The vector relationships are expressed from the outset in novel environments with novel objects. Object-vector cells are distinct from grid cells, which use a distal reference frame, but the cells exhibit some mixed selectivity with head-direction and border cells. Collectively, these observations show that object locations are integrated in metric representations of self-location, with specific subsets of MEC neurons encoding vector relationships to individual objects.

## Introduction

The hippocampus and the medial entorhinal cortex are part of a brain system for mapping of self-location that is likely recruited when animals navigate between locations in the proximal environment^1–5^. In the hippocampus, place cells fire specifically when the animal is at certain places^6,7^, and goal-vector cells encode the animal’s movement in the direction of a goal^8,9^. Collectively, intermixed assemblies of place cells and goal-vector cells may create maps of the animal’s present and future position in the local environment^1,10^. Each individual environment has a unique map^11–14^ that when combined with information about events in the environment^15^ and their temporal order^16^ forms the basis for location and route-based episodic memories^17,18^. Hippocampal place and goal representations are likely to interact with spatial maps in the adjacent medial entorhinal cortex (MEC)^2,19,20^. There position is encoded by multiple cell types, including grid cells, which fire at periodic locations that produce a hexagonal array across the entire available space^21^, border cells, which fire if and only if animals are near the geometric borders of the environment^22,23^, and an independently regulated class of cells with discrete but irregularly patterned spatial firing fields^24^. These location-coding neurons are supported by cells that by encoding head direction^25,26^ and speed^27^ may allow the representation of position to change instantaneously and dynamically in accordance with the animal’s movement in the environment, independently of environmental context^28–31^. As a whole, this circuit of space-coding neurons in hippocampus and entorhinal cortex is thought to form a key element of the braińs neural network for goal-directed spatial navigation^1,2^.

While the number of spatial circuit elements is unfolding at an accelerating rate^5^, insights are limited by their almost exclusive reliance on recordings from rodents foraging in empty enclosures very different from the richly populated, geometrically irregular environments of the native world^3,4^. Natural environments contain objects, and animals may use these for navigation. In the lateral division of the entorhinal cortex (LEC), a subset of cells fire specifically when animals encounter discrete objects in the recording environment, and firing fields at the locations of these objects are often maintained even after the objects are removed^33,34^. But how representations of object locations are used to navigate in the space between objects remains to be determined.

Behavioural studies have shown that rodents^35–38^, as well as birds^39^ and insects^40^, store information about distance and direction to discrete landmarks in the environment, and that this allocentric vector information can be used to guide navigation. Theoretical studies have further proposed that landmark-dependent vector operations give rise, in the mammalian brain, to cells with firing fields that depend on distance and direction from discrete objects^41^. Other theoretical work has suggested the existence of ‘boundary vector cells’ with firing fields tuned specifically to distance and direction from walls, and other extended boundaries, rather than from confined objects^42–44^. Evidence for vector representations of either kind is limited. One study identified a small number of hippocampal CA1 cells (less than 10) that fired at distinct directions and distances from discrete objects, consistent with a landmark-vector encoding operation^45^. The firing fields of these cells developed only at certain objects and mostly over a time course of multiple trials, suggesting that these cells cannot alone subserve landmark-based navigation, which should operate almost instantly. Other studies have reported cells that encode vectors to wall-shaped boundaries^46,47^ but the majority of these cells fire not primarily in the open space of the arena but mostly alongside the boundaries^47^, mirroring the firing fields of border cells in the MEC^23,48^. The ability of these cells to encode vectors to all possible positions is further limited by the fact that there are often few straight boundaries in the animal’s natural habitat. Vector navigation would therefore benefit a lot if cells were able to use also point-like landmarks as references. In the present study, we looked for such cells. Our search focused on the superficial layers of MEC, the very same brain network that by way of grid cells represents position on content-free planar surfaces. We report that when discrete objects are present in the environment, the animal’s direction and distance from the objects is encoded in the activity of a substantial number of MEC cells, comparable to the number of grid cells in the same brain region.

## Results

### Object-vector cells in the medial entorhinal cortex

We recorded neural activity from a total of 503 cells in 8 mice. Post-hoc histological analyses showed that the tetrode tracks terminated in the superficial layers of MEC (Extended Data Fig. 1). The mice foraged freely for cookie crumbs in a compartment with a floor that was either square (0.8×0.8×0.5 m or 1.0×1.0×0.5 m, 247 cells) or circular (diameter: 0.9 m or 0.65 m, height: 0.5 m, 256 cells). A typical experiment began with a trial where no object was present in the compartment. This trial was succeeded by a trial where a tower-like object was placed on the floor of the compartment before the start of the trial (Fig. 1a).

**Figure 1.**
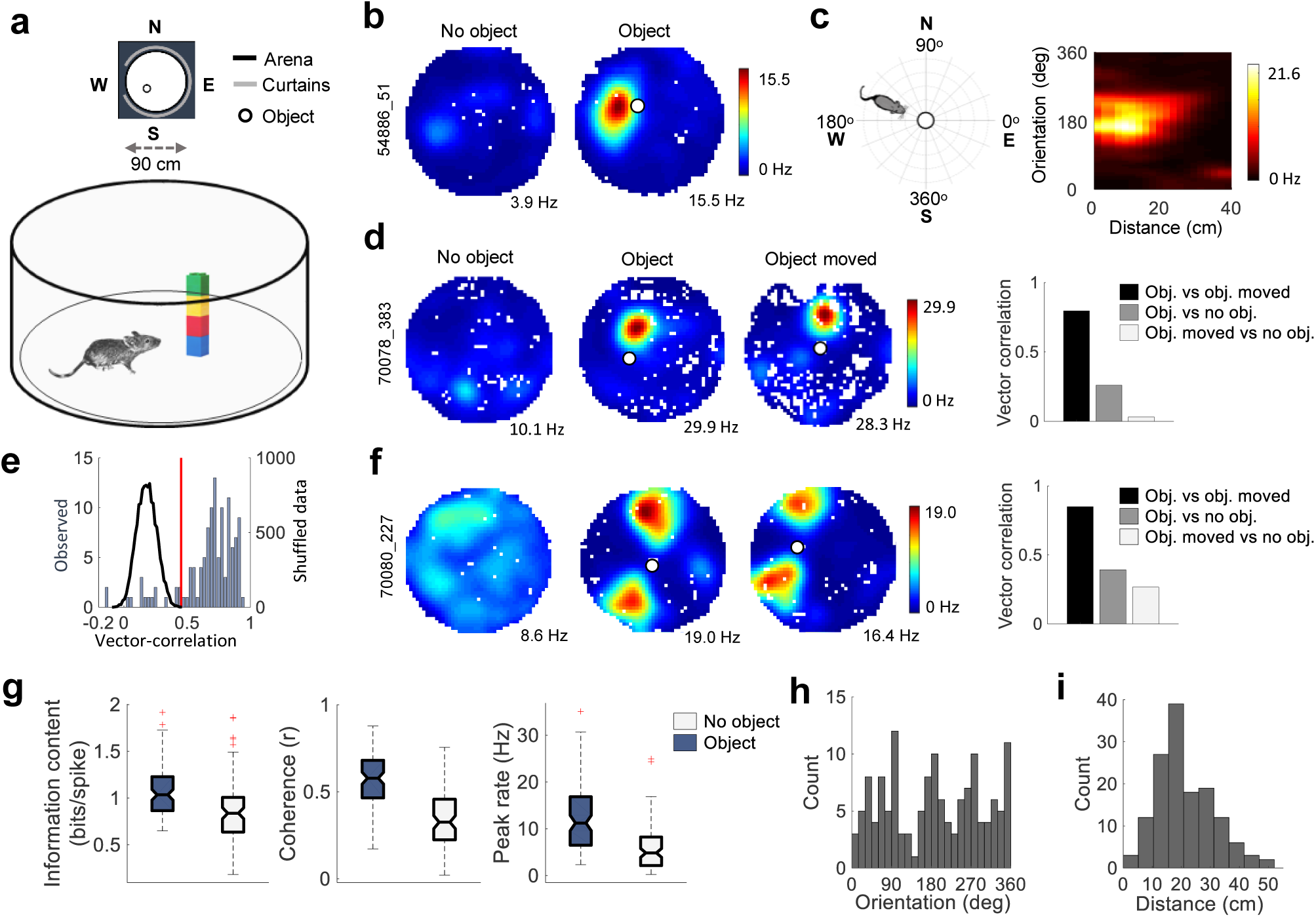
Object-vector cells fire at fixed distances and directions from an object. **a)** Upper panel: schematics of recording environment (birds-eye view). Object location (small circle) varied between trials. Lower panel: mouse exploring arena with object. **b)** Colour-coded firing rate maps showing example cell that expressed a new firing field when an object (white circle) was introduced into the recording enclosure (right). Rate map for trial before introduction of object is shown to the left. Peak rates are indicated below rate maps. Mouse and cell numbers are indicated in vertical text bar (5 and 2 digits, respectively). **c)** Left: Firing rates were compared across angular bins of 5 degrees and distance bins of 2 cm, with the centre of the object as a reference for both direction and distance. Zero degrees is defined as east (in the room frame). Right: Colour-coded vector map showing firing rate for the example cell in b as a function of distance (*x*-axis) and orientation (*y*-axis) from the object. **d)** Left panels: Colour-coded rate maps for a different example cell on three successive trials (peak rates below rate maps; mouse and cell number in vertical text bar). The first trial was conducted without an object in the arena, the second with an object, and the third with the object in a different position. Right: correlation between vector-maps (generated as in c) for the three pairs of trials (same cell as to the left). **e)** Frequency distribution showing correlations between vector maps on trial pairs with objects in different positions (second and third trial in d). Blue bars: trial-pair correlations of the 120 cells that expressed new fields in the presence of the object (counts on left *y*-axis). Black line: Correlations in shuffled versions of the same data (counts on right *y*-axis)). Red line marks 99th percentile of distribution of shuffled data. **f)** A subset of object-vector cells had two object-vector fields. Left: rate map for one such cell over three trials as in d. Object location is indicated by white circle. Right: correlation between vector maps (same cell as in the rate maps). **g)** Box plots showing spatial information content (bits/spike), spatial coherence (correlation between adjacent bins), and peak firing rate for all object-vector cells on the no-object trial and the first trial with the object. In this and subsequent box plots, horizontal black lines indicate median values, box edges indicate 25^th^ and 75^th^ percentiles, whiskers extend to the most extreme point that lies within 1.5 times the interquartile range, and data points larger than 1.5 times the interquartile range are shown as outliers (red crosses). **h)** Frequency distribution showing peak orientation of all fields in the vector-map that were stable between object-and displaced-object trials. **i)** Frequency distribution showing peak distance of all fields in the vector-map that were stable between object and displaced-object trials.

We first asked if the activity of any of the recorded MEC cells was changed by the very presence of an object. For this experiment we generally used a 5 cm wide, 20 cm tall cylinder-shaped tower (Object 7, Extended Data Fig. 2). Specifically, we looked for object-induced firing fields. Firing fields were defined by the stepwise contour surrounding areas with firing rates ranging from two times the standard deviation across bins of the rate map to the peak rate of the map. Contiguous areas within a contour of at least 16 bins and with a peak firing rate of at least 2 Hz were counted as firing fields. On trials with the object in the recording enclosure, 237 cells showed significant spatial information, exceeding the 95th percentile of a distribution of spatial information values for shuffled versions of the data (0.65 bits/spike). Among these cells, 120 had firing fields that were not present when the same cells were recorded in the absence of the object. In general, the centre of these fields was offset from the object by several centimetres, often several tens of centimetres. Only 3 of the cells (from 2 animals) had firing fields at the location of the object (< 4 cm from centre of field to centre of object). In the majority of cells that had displaced firing fields (offsets of more than 4 cm), the vector relationship (distance and direction from the object) was maintained when the object was moved to a new location on a subsequent trial (Fig. 1d), suggesting that the cell’s activity was determined by the location of the object and not the distal cues.

For the 117 cells that exhibited displaced firing fields, we quantified the relationship between spatial firing and the object by constructing vector maps that expressed firing rate as a function of both direction and distance to the centre of the object (Fig. 1c). A cell was categorized as an object-vector cell if the correlation of the vector maps from the two trials with objects at different locations exceeded (i) the 99^th^ percentile of a distribution of correlations between rate maps where spike times had been randomly jittered (99^th^ percentile correlation value: 0.45) and (ii) the correlations between each object trial and the no-object trial. A total of 98 out of the 117 cells passed these criteria (Fig. 1e). Among these object-vector cells, as expected, the vector from object to field centre on the first object trial predicted the location of a firing field on the second trial (Fig. 1d; Extended Data Fig.3). Fifty-five of the cells (56%) had only one object-induced firing field; 42 (43%) had two fields (Fig. 1f), and 1 cell had three. Cells with two or more fields were not grid cells as they consistently had grid scores below chance levels and no firing was observed in regions defined by a hexagonal matrix extrapolated from the two or three identified fields (Extended Data Fig. 4). The presence of multiple fields was not caused by poor cluster isolation; cells with two fields were as well-separated as cells with one field (isolation distances of 44.9 ± 5.3 and 44.2 ± 4.3, respectively, means ± s.e.m.; Extended Data Fig. 5). Object-vector cells had a higher spatial information content (Wilcoxon signed rank test, W = 4024, n1 = n2 = 98, P = 1.5×10^−8^), a higher spatial coherence (W = 4753, n1 = n2 = 98, P = 1.6×10^−16^), and a higher peak firing rate (W = 4616, n1 = n2 = 98, P = 8.3×10^−15^), on object trials than on no-object trials (Fig. 1g).

We determined a field’s orientation and distance from the object by identifying the peak of the firing field in the object-referenced vector-map. The orientations of object-vector fields from different cells covered the entire azimuthal range (Fig. 1h). The directional preferences of the fields were similar in square and circular compartments, suggesting they were independent of the internal geometry of the box (Extended Data Fig.6). For cells with multiple fields, the mean angular difference between pairs of fields was 131.5 ± 4.3 degrees (42 cells; Extended Data Fig. 7a,b; mean ± S.E.M.). The smallest angular difference was 75 degrees. The distribution of field-distances from the object showed a skew towards proximal locations, with a mean centre-to-centre distance of 21.2 ± 0.8 cm (Fig. 1h, mean ± s.e.m.), clearly distinguishing these cells from the object-centered cells of the LEC^33,34^ and the very few cells with such properties in the present MEC data. The largest field distance among the object-vector cells was 52 cm. Distances between the centre of object and the contour of the field ranged from 0 cm to 33 cm (mean ± s.e.m.: 8.9 ± 0.7 cm). Field size was not significantly correlated with distance from the object (Extended Data Fig. 7c; r = 0.15, P = 0.07).

### Object-vector cells are allocentric

Object-vector cells were defined as cells that fired at specific locations in space regardless of the animal’s own orientation. To test whether object-vector cells encoded vectorial information also in an egocentric reference frame, we constructed egocentric tuning curves (heading direction vs. firing rate) for each cell by counting spikes as a function of movement direction relative to the object, using procedures similar to those employed to identify egocentric goal-vector cells^9^ (18 bins of 20° each; time was normalized to time spent moving in each direction relative to the object). We defined 0° as moving towards the object and ± 180° as moving away from the object (Fig. 2a). In general, object-vector cells exhibited no egocentric tuning to the object (Fig. 2 b-d). Cells emitted similar numbers of spikes within allocentrically defined fields even when the heading directions relative to the object were opposite (Fig. 2b,c). To quantify the tuning to movement direction relative to the object, we defined an egocentric directionality index as the difference between the peak firing rate and the median rate of the movement-direction tuning curve, divided by the curve’s median absolute deviation. Only 3/98 object-vector cells, no more than expected by chance, had an egocentric directionality index that exceeded the 99^th^ percentile of a distribution of directionality indices obtained from shuffled versions of the data (Fig. 2 d,e). The median value for the index was 2.25 (50^th^ and 99^th^ percentiles of shuffled data: 2.3 and 9.2, respectively). Object-vector cells thus encode directions and distances to landmarks independently of the animal’s heading movement towards the object, as expected if the object vectors of the recorded MEC cells are coded exclusively in an allocentric framework.

**Figure 2.**
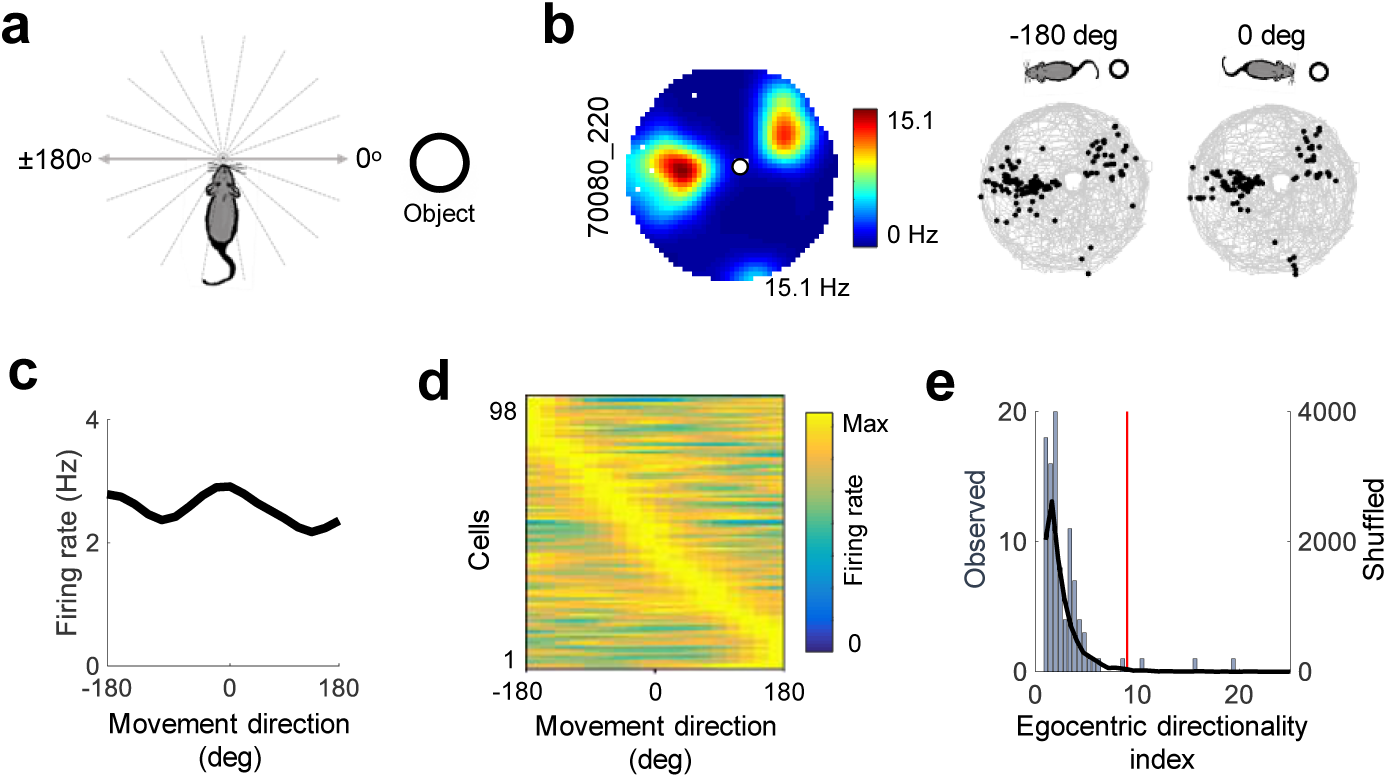
Object-vector cells are allocentric. **a)** Egocentric reference frame. Zero degrees is defined as moving towards the object (white circle), 180 degrees as moving away from the object. **b)** Left: Colour-coded firing rate map for a representative object-vector cell (peak rate below rate map; mouse and cell number indicated in vertical text bar). Right: path plot showing, for the same cell as in the rate map, the animal’s path with overlaid spike locations in black. Left path plot shows spikes on trajectories away from the object; right plot shows spikes on trajectories towards the object. **c)** Egocentric directional tuning curve for the cell in b. Firing rate is shown as a function of direction of movement relative to the object. Directional bins were 20 degrees. The curves were smoothed over 1.5 bins (30 degrees) with a Gaussian filter. Note that egocentric directional modulation is nearly absent. **d)** Colour-coded egocentric directional tuning curves (as in c) for all object-vector cells. Each horizontal line corresponds to one cell and shows firing rate colour-coded as a function of movement direction. Cells are sorted according to the movement direction that had the highest firing rate (light yellow). Note the relative absence of egocentric directional tuning. **e)** Distribution of egocentric directional modulation across the entire sample of object-vector cells. Egocentric directional modulation was estimated by defining for each cell an egocentric directionality index as the difference between peak firing rate and median firing rate in the egocentric tuning curve (plotted as in c), divided by the median absolute deviation. Distribution of observed values is shown as blue bars (counts on the left *y*-axis), shuffled data as black line (counts on the right *y*-axis). Red line marks the 99th percentile of egocentric directional modulation values for the shuffled data.

### Object-vector cells generalize between objects

To determine whether object-vector cells respond to specific objects or represent directions and distances from any object, we recorded 22 of these cells from 6 mice in experiments where the number of simultaneously presented objects was increased to between 2 and 6. Objects were selected from a pool of tower-shaped objects. Some of these were fully or partly shaped like rectangular prisms, others had a more cylindrical appearance (Extended Data Fig. 2). In total, we tested 49 object-cell combinations (2-3 objects for most of the 22 cells). In 48 of the combinations, new firing fields emerged when the object was introduced in the arena (Fig. 3a). The distance and direction relationship to the object was largely retained from one object to the other, i.e. there were only small differences between object pairs in distance and direction from the nearest object (Fig. 3b; distance difference: 3.7 ± 0.5 cm; direction difference: 7.7 ± 1.0 degrees; means ± s.e.m.). These relationships were maintained even in experiments where objects were flattened to nearly two dimensions by shrinking either the width or the height of the object to less than 0.5 cm (Objects 6 and 13, respectively, in Extended Data Fig. 2; Extended Data Fig. 8a,b).

**Figure 3.**
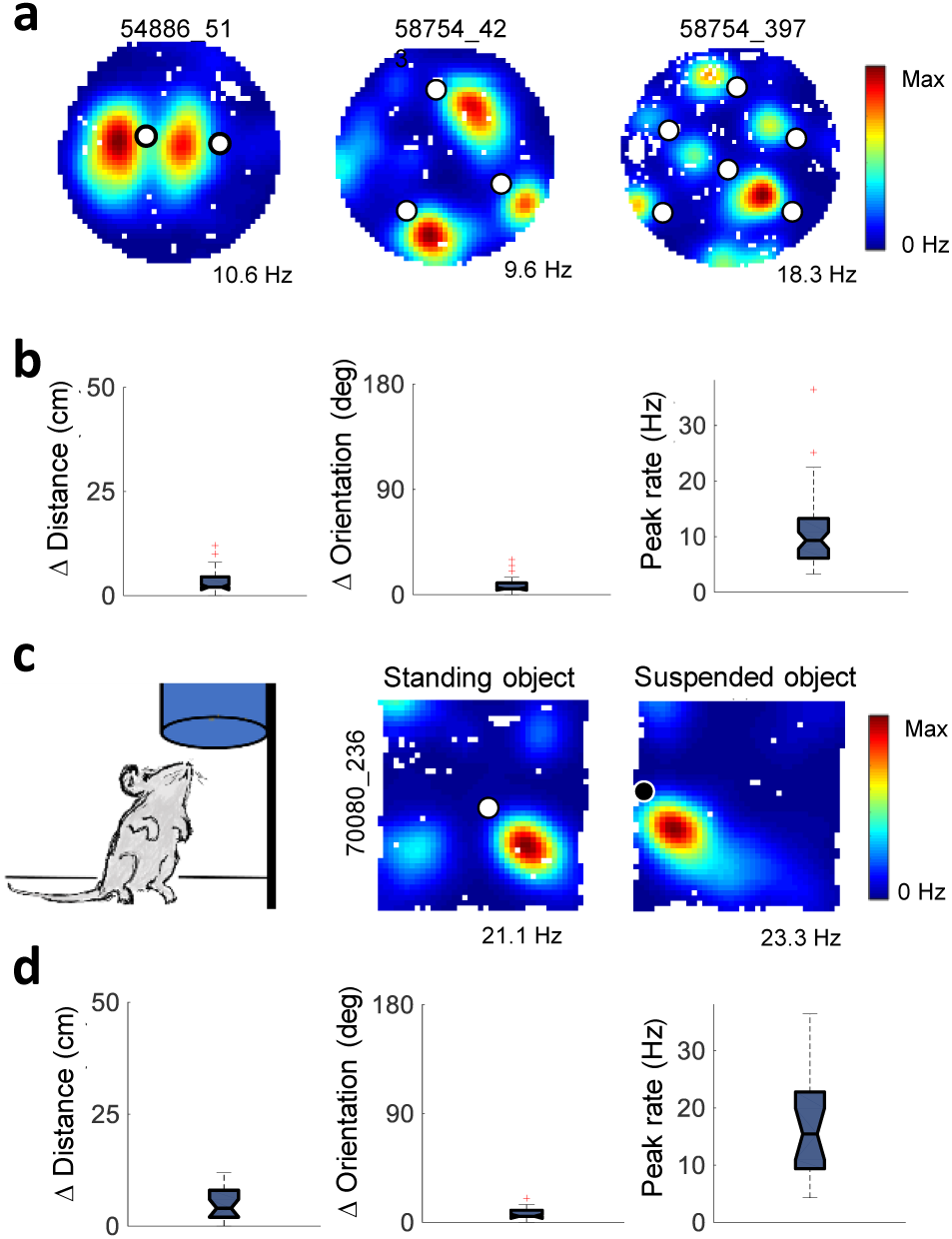
Object-vector cells generalize between objects. **a)** Colour-coded firing rate maps showing three cells recorded with two, three, or six objects (white circles) placed at the same time in the arena. Mouse and cell number at the top, peak rate at the bottom. **b)** Box plots showing, for all pairs of object-vector fields recorded in the same multi-object experiment, the difference in peak distance from object (left) and the difference in orientation of field relative to object (middle). The differences were calculated from the nearest pair of fields in the vector-maps made for each object location. Right: peak firing rates across all firing fields on multi-object trials. **c)** Left: mouse exploring suspended object attached to a wall of the recording box. Right: colour-coded firing rate maps showing example cell recorded successively with standing object (white circle) and suspended object (black circle). Mouse and cell number in vertical text bar; peak rate at the bottom. Note similar vector tuning for standing and suspended objects. **d)** Box plots showing, for all object-vector cells tested with a suspended object, the difference in distance between field and standing vs. suspended object (left) and the difference in orientation of field relative to standing vs. suspended object (middle). Differences were calculated as in b. Right: peak firing rates of object-vector fields elicited by suspended objects.

We next asked whether object-vector cells would differentiate between objects that obstructed the path of the animal and objects that were not functional barriers but still stood out from the visual background. To test this, a cylinder-shaped object was attached to one of the walls of the recording box, with the bottom of the object 15 cm above the ground level (Fig. 3c). The mice moved freely underneath the suspended object and could not reach it from an upright position. A total of 21 object-vector cells were recorded from 5 mice tested with both standing and suspended objects. In all of these cells, a new firing field was expressed in response to the lifted object (Fig. 3c). As in experiments with multiple standing objects, the vector relationship was retained across the pair of objects (Fig. 3 c,d; difference in horizontal distance: 4.0 ± 0.8 cm; difference in direction: 7.7 ± 1.3 degrees, means ± s.e.m.), suggesting that object-vector fields emerge regardless of whether the object interferes with the animal’s path or not.

The fact that also suspended objects generated object-vector fields points to visual input as one likely source of location-specific firing in these cells. We tested this further by comparing rate maps of object-vector cells in the presence of an object before and after the room lights were turned off while the animal was in the box (21 object-vector cells from 5 mice). Correlations between vector maps on light and dark trials were significantly decreased compared to correlations for pairs of light trials in the same cells (Extended Data Fig. 9ab; Wilcoxon signed rank test, W = 207, n = 21, P = 0.0015). Spatial information and spatial coherence were also reduced (Extended Data Fig. 9c). For some cells, neither of these measures were reduced to chance levels, however, possibly reflecting the continued use of path integration in darkness.

### Object-vector responses do not require experience

We then asked whether object-vector cells require temporally extended or repeated experience with the object or the recording environment to encode vectors from the object to locations in the open space. We recorded 19 object-vector cells in two different rooms, A and B, distributed over 6 experiments in 4 mice. Thirteen of these cells were recorded during the animal’s first exposure to room B, and the objects used during this encounter were all unfamiliar to the animal. The objects were selected from the pool of objects in Extended Data Fig. 2. All cells with object-vector properties in the familiar room (room A) had object-vector fields also in the new environment with new objects (room B), and there was no significant difference in the spatial information content of the rate maps in the two settings (Fig. 4a,b; familiar: 1.10 ± 0.05 bits/spike; novel: 0.96 ± 0.09 bits/spike, means ± s.e.m.; Wilcoxon signed rank test, W = 66, n = 13, P = 0.17), suggesting that object-vector cells were sharply tuned from the first trial when the objects were encountered. When we realigned vector maps to compensate for the reorientation that typically occurs in MEC cells when animals move from one environment to another^23,28,29^, correlations between vector maps in familiar and new rooms became very high (Fig. 4c; r = 0.81 ± 0.02, mean ± s.e.m.), suggesting that individual cells maintained their distance metrics.

**Figure 4.**
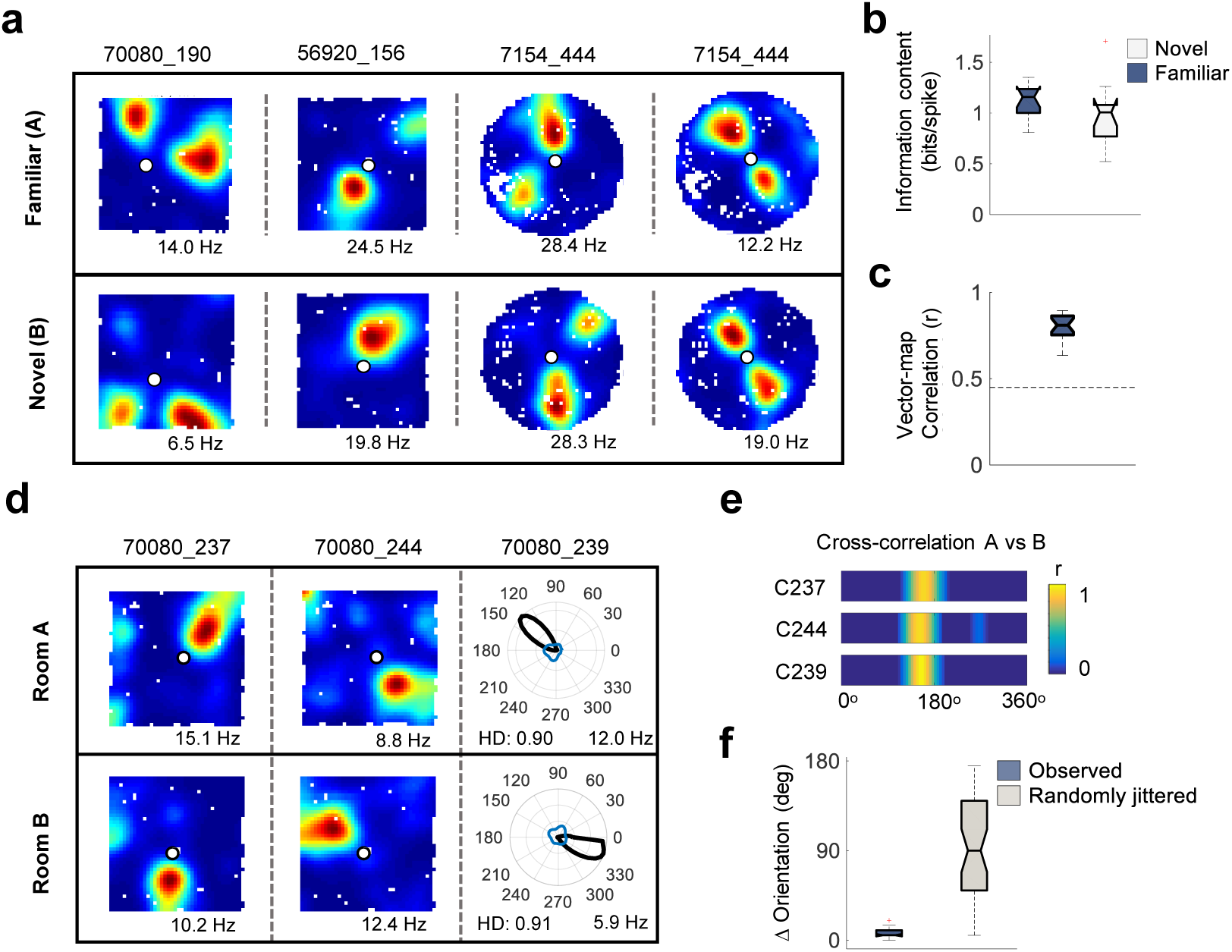
Object-vector fields with novel objects in novel environments. **a)** Colour-coded rate maps for four example cells recorded in both a familiar environment and a novel environment with a novel object (rooms A and B; top and bottom rows, respectively). White circles represent objects. Mouse and cell numbers at the top; peak rates at the bottom. **b)** Box plot showing spatial information content for the 13 cells recorded successively in familiar and novel environments. **c)** Box plot showing vector-map correlation between rooms A and B for all 13 cells recorded where room B was novel and for all 6 additional cells recorded when both rooms had been experienced before. Vector maps were constructed as in Fig. 1c. Vector-maps were shifted circularly with respect to orientation until maximal correlation was obtained, in order to compensate for orientation realignment between rooms. High correlations imply that pairs of rate maps in rooms A and B were essentially rotations of each other. **d)** Rate maps for two object-vector cells and one head direction cell recorded concurrently in room A (top) and room B (bottom). Symbols as in a. Both rooms were familiar in this experiment. **e)** Color-coded cross-correlations of directional tuning curves in room A and B for the 3 object-vector cells in d. Cell number is indicated on the left for each color-coded cross-correlation. Note maximal correlation near an orientation shift of approximately 160 degrees for all three cells. **f)** Box plot showing difference in orientation at peak cross-correlation between rooms A and B for all pairs of cells recorded in both rooms, compared to differences in peak cross-correlation for the same cell pairs after randomly shifting the orientation tuning curves of the cells.

In 5 of the 6 experiments, we recorded more than one object-vector cell, and object-vector cells were also recorded along with other spatially or directionally modulated cells. In total, there were 50 pairs of simultaneously recorded cells in experiments with two recording rooms (18 object-vector cells, 4 head-direction cells, 1 grid cell). In order to compare shifts in the directional preferences of object-vector cells and head-direction cells between rooms, we calculated circular cross-correlations between directional or spatial tuning curves for the two rooms. For each cell (object vector cell, head direction cell, or grid cell), the shift in orientation between rooms was defined as the rotation of the tuning curve or rate map from one room that maximized the correlation with the tuning curve or rate map in the other room. Then, for each pair of simultaneously recorded cells (object-vector cells or other cells), we determined the difference in angular shift between the two rooms for the two cells. The distribution of pairwise rotation differences was then compared with a distribution obtained from pairs of cells in randomly jittered versions of the tuning curves. The observed pairwise differences were significantly smaller than differences between randomly shifted tuning curves, both for pairs of simultaneously recorded object-vector cells (observed: 8.2 ± 0.8 degrees for all cell types; random: 88.8 ± 7.4 degrees, means ± s.e.m.; Wilcoxon signed rank test, W = 10, n = 50, P = 2.0×10^−9^) and for pairs consisting of one object-vector cell and either one head-direction or one grid cell (observed: 10.3 ± 1.3 degrees for all cell types; random: 92.8 ± 13.0 degrees, means ± s.e.m.; Wilcoxon signed rank test, W = 1, n = 16, P = 5.3×10^−4^) (Fig. 4 d-f). Grid cells and head-direction cells generally maintained stable orientations across object and no-object trials (Extended data 10a,b; spatial correlation of grid cells between object and no-object trials: 0.63 ± 0.04, mean ± s.e.m.; correlation of head-direction tuning curves between object and no-object trials: 0.54 ± 0.04).

These observations suggest that a fixed directional structure was maintained across all cell types from one environment to the other. The fact that object-vector cells rotated coherently with grid cells and head direction cells in the two-room experiments raises the possibility that object-vectors cells bind local cues to the global distal framework that controls head direction cells and grid cells.

### Object-vector cells are distinct from grid cells and head direction cells

We next asked if object-vector cells constitute a subpopulation of previously reported spatially-modulated MEC cells, or if they represent a new and separate functional population. Out of the 503 superficial MEC cells recorded in the present study, 40 cells, or 8.0%, were identified as grid cells, 57 (11.3%) qualified as speed cells, 149 (29.6%) as head-direction cells, and 34 (6.8%) as border cells.

The majority of the 98 object-vector cells (19.5% of the 503 cells) generally did not share defining properties with any other spatially or directionally modulated cell types (Fig. 5ab). First, they were clearly distinguishable from grid cells (Fig. 5ab; Extended Data Fig. 10a). A significant object-vector response was observed only in one grid cell (1.0%), a number not significantly different from numbers expected with random selection from a shuffled distribution (binomial test with expected P_0_ of 0.01, P(X≥1) = 0.63; P(X≤1) = 0.74), but significantly lower than the number of cells expected to pass criteria for both grid cells and object-vector cells with the currently identified numbers of these cells, if these cell populations were independent (expected number = N(fraction of grid cells × fraction of object-vector cells) = 7.8; binomial test, P(X ≤ 1) = 0.003). Consistent with the low number of grid cells with object-vector tuning, the grid scores of the object-vector cells, assessed in the no-object condition, were significantly lower than in all other cells (Fig. 5b; Mann-Whitney U-test, U = 19949, n1 = 405, n2 = 98, P = 0.0018). Second, object-vector cells were distinct from speed-tuned cells. The number of object-vector cells that also passed the criterion for speed cells on no-object trials was 5 (5.1%) – more than expected with random selection from a shuffled distribution (binomial test with expected P_0_ of 0.01, P(X≥5) = 0.003), but lower than the number of object-vector cells expected to be speed-modulated based on the overall level of speed modulation in the general MEC population (expected number, calculated as for overlap with grid cells: 11.1; binomial test, P(X≤= 0.04). There was no difference in speed scores of object-vector cells compared to the other cells recorded (Fig. 5b; U = 23369, n1 = 405, n2 = 98, P = 0.29). Third, and in contrast to the limited overlap with grid cells and speed cells, as many as 38 (38.8%) of the object-vector cells passed the criterion for head-direction cells on the no-object trial (Fig. 5 a,b; Extended Data Fig. 10b). This number was larger than expected with random selection from the distribution of shuffled data (binomial test with expected P_0_ of 0.01, P(X≥38) = 1×10^−8^), but not significantly higher than the number expected to pass both head-direction criteria and object-vector cell criteria based on the number of head direction cells in the overall MEC population, if the two cell populations were independent (expected number: 29; binomial test, P(X≥38): 0.06). In the majority of object-vector cells that passed the head-direction criterion, the head-direction mean-vector-length scores were low (mean ± s.e.m.: 0.38 ± 0.02; pure head direction cells: 0.59 ± 0.02, Mann-Whitney U-test, U = 1.9×10^3^, n1 = 38, n2 = 114, P = 1.4×10^−5^), yet the head-direction scores, estimated on no-object trials, were significantly higher for object-vector cells than for the remaining MEC cells (mean score ± s.e.m.: 0.24 ± 0.015; U = 29527, n1 = 405, n2 = 98, P = 2.2×10^−4^). Taken together, these data suggest that a fraction of the object-vector cells were weakly head direction-tuned during free running in the open arena.

**Figure 5.**
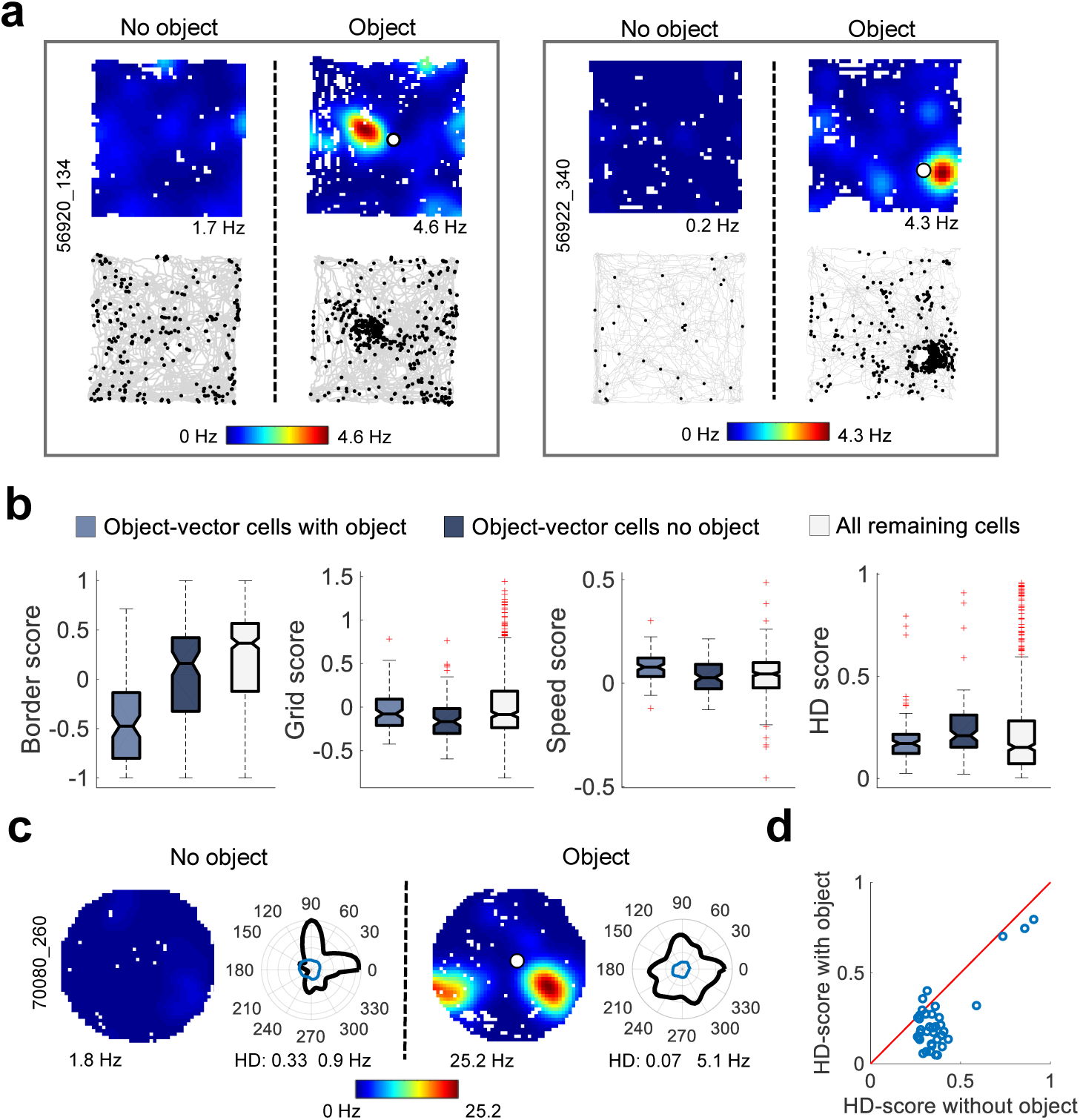
Object-vector cells are distinct from other spatially-modulated cell classes. **a)** Colour-coded rate maps (top) and path plots (bottom) for two representative object-vector cells recorded first without an object and then with an object (white circle). Path plots show the animal’s trajectory with spike locations superimposed as black dots. Peak rates and mouse and cell number are indicated (horizontal and vertical text labels). The cells responded vigorously to objects but did not have border or grid fields. **b)** Box plots showing distribution of border scores, grid scores, speed scores, and head-direction (HD) scores for object-vector cells recorded without an object (dark blue boxes) or with an object (light blue boxes), and for cells that did not satisfy criteria for object-vector cells on the no-object trial, including all other types of spatially or directionally modulated cells (light grey boxes). Note the presence of outliers in the distribution of head direction scores in object-vector cells, suggesting that a subset of object-vector cells is modulated by head-direction input. **c)** Response to object in an object-vector cell tuned to head direction of the mouse. Left: recording with no object (left) and with an object (right). For each trial, a colour-coded firing rate map is shown along with a circular plot for firing rate as a function of head direction (black curve, firing rate; blue curve, time spent; HD, head direction score). Peak rates are indicated for rate maps as well as circular plots. Note that head direction tuning (HD) is reduced when the object is present. **d)** Head-direction (HD) score for all object-vector cells that passed the head-direction criteria on the no-object trial, plotted against the head-direction score of the same cells on the object trial.

We then asked how tuning by head direction was influenced by introduction of an object on the floor of the arena. In general, the object decreased the tuning to head direction (Fig. 5b-d; Extended Data Fig. 10b). The number of cells passing the criterion for head direction cells on the object trial was reduced to 20 (20.4% of the cells). While this was still more than expected with random selection from shuffled data (binomial test with expected P_0_ of 0.01, P(X≥20) = 1×10^−8^), it was significantly lower than the number of cells expected to share head-direction and object-vector properties in the entire population of recorded cells if these two cell types were independent (binomial test, expected number: 29, P(X≤20) = 0.04). Similarly, head direction scores for object-vector cells were significantly reduced on object trials compared to no-object trials (W = 3555, n = 98, P = 6.3×10^−5^; head direction scores for object-vector cells vs. remaining cell population on the object trial: U = 25827, n1 = 405, n2 = 98, P = 0.40). There was no difference in the distribution of time spent across the spectrum of head directions on object and no-object trials (Fig. 5c; Extended Data Fig. 10b; mean vector lengths of head-direction occupancy, Wilcoxon signed rank test, W = 1967, n = 98, P = 0.10). The loss of head-direction tuning on object trials could be taken to imply that head direction inputs exert a strong influence on firing in object-vector cells when objects are not present, whereas upon object insertion they combine with inputs from cells that signal object location^33,34^ or the animal’s egocentric orientation to the object^9,49,50^ to form an allocentric object-vector representation, much in the same way as head direction inputs dominate firing in grid cells when hexagonally patterned firing breaks down after blockade of hippocampal excitatory feedback^51^. Taken together, these observations and those from the two-room experiment suggest that object-vector cells are modulated by head direction inputs, but this modulation may be secondary to influences from object-informative cells and does not account for the presence of a distinct population of cells expressing vectorial relationships to local objects.

### Object-vector cells are distinguishable from border cells

Next, we asked to what extent object-vector cells overlap with border cells, since objects and borders can be seen as overlapping and potentially continuous stimulus populations. The data suggest that object-vector cells and border cells exhibit only partial overlap. Most object-vector cells lacked firing fields along the peripheral walls of the recording enclosure (Fig. 5a), and border scores on the no-object-trial were lower for object-vector cells than for cells that were not object-vector cells (Fig. 5b; Mann-Whitney U-test, U = 20680, n1 = 405, n2 = 98, P = 0.0017). Yet, among the 98 object-vector cells recorded on the no-object trial, as many as 12 passed the criterion for border cells (35.3% of the border cells; 12.2% of the object-vector cells; Fig. 6 a,b). The number of object-vector cells that also passed the border cell criteria was larger than expected with random selection from a shuffled distribution (binomial test with expected P_0_ of 0.01, P(X≥11) = 1×10^−8^), and larger than the number expected if the properties of border cells and object-vector cells were independently distributed in the MEC population (expected number: 6.6; binomial test, P(X≥12) = 0.04).

**Figure 6.**
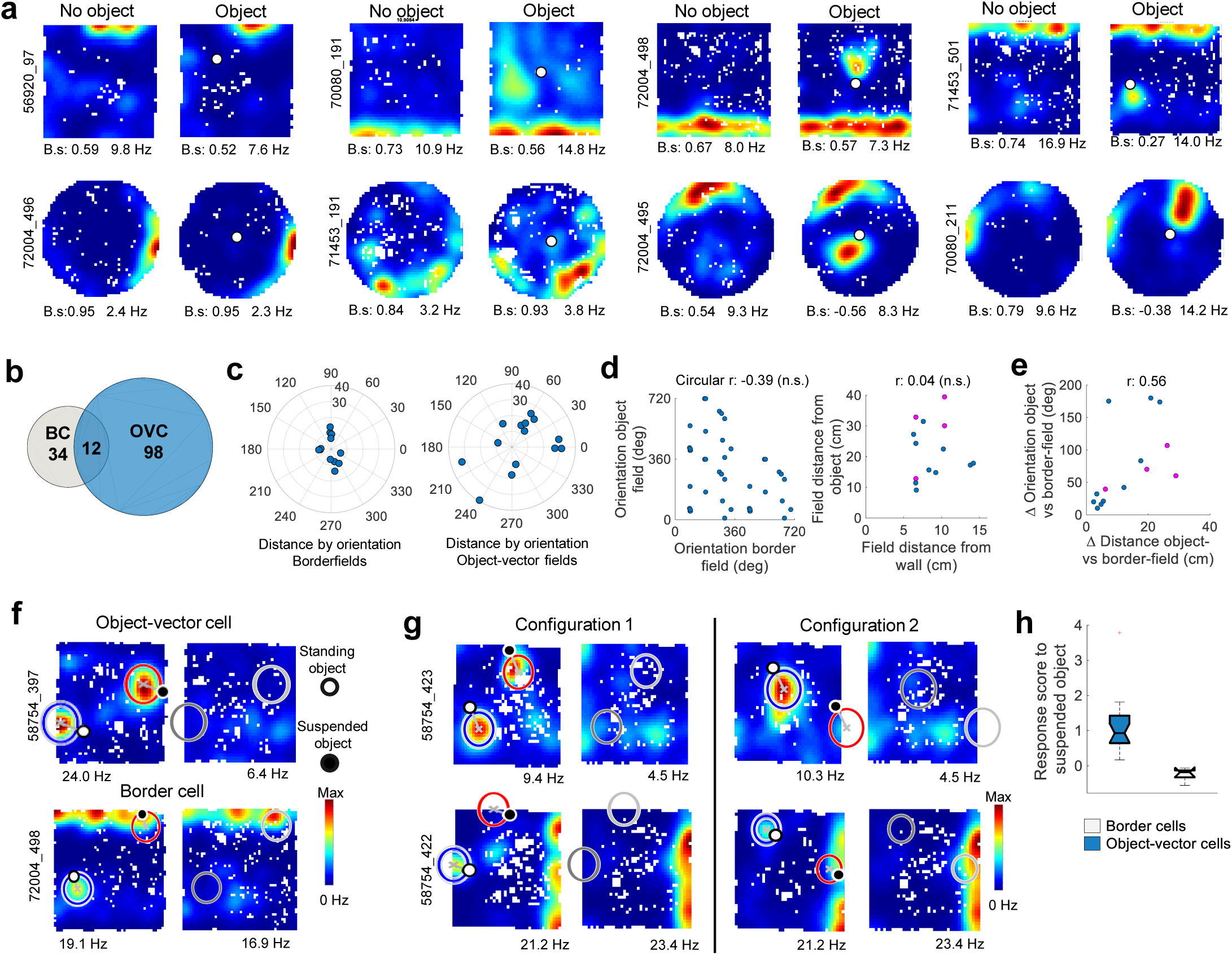
Object-vector cells overlap with a subset of bordercells. **a)** Color-coded rate maps of 8 border cells recorded with and without an object (white circles), illustrating a continuum of object-responsiveness in border cells. The two border cells on the left – one recorded in a square and one in a circle – show no response to the object, the two cells in the second column show weak changes between object and no-object trials, and the four cells to the two right columns display clear firing fields in response to the object. **b)** Venn diagram showing overlap between populations of object-vector cells (OVC) and border cells (BC). Twelve out of 34 border cells also passed the criteria for object-vector cells. **c)** Lack of relationship between vectorial tuning to border and object for cells that responded to both border and object. Left: radial plot showing distribution of border-field vectors. Right. The same for object-field vectors of cells that responded to both borders and objects. Length of units (10 cm) corresponds to distance of field centre from object or border (same scale for both diagrams). Note wide distribution of orientations and distances for object-vector fields but less so for border fields. **d)** Relationship between vectors to wall and object for cells with tuning to both. Left: lack of correlation between orientations of border-fields and object-fields. Because of the circular nature of orientation, the orientation of the object-field vectors and the border-field vectors are plotted over two cycles, i.e. each data point is shown twice. A correlation between orientations to border and object would be reflected as regression towards the diagonal and parallel lines offset by 360 degrees from the diagonal. Right: lack of linear correlation between distance from wall to border-field and distance from object to centre of object-vector field. Note that the distance to the object is consistently larger than to the border. Purple dots mark pairs of cells that expressed two object-vector fields but only one border field. **e)** Difference in orientation of border-field vector and object-field vector plotted against difference in length of vector between wall and centre of border-field and between object and centre of object-field. Differences were significantly correlated, i.e cells that differed with respect to orientation also differed with respect to orientation. Purple circles as in e. **f)** Procedure for comparing response to standing and suspended objects in colour-coded rate maps. Area and vector defined by response to the standing object is transferred to the location of the suspended object to act as a template for identifying a possible corresponding firing field there. Mean firing rates were determined within both of these areas in the presence and absence of objects. Standing and suspended objects are shown as black and white circles, respectively. Left: object-vector cell; right: border cell. Note that in the object-vector cell, firing rate is increased similarly at the standing and the suspended object, whereas in the border cell there was no enhancement of firing near the suspended object. **g)** Colour-coded rate maps showing one object-vector and one border cell recorded concurrently in the presence of both a standing and a suspended object, over two trials where object locations were changed. White circle, standing object; black circle, suspended object. Peak rates (horizontal) and mouse and cell numbers (vertical) are indicated. **h)** Box plot showing differential response of object-vector and border cells (entire sample of 20 object-vector cells and 5 border cells). Responses to suspended objects were defined by calculating a score expressing the difference between the firing rate in the template area near the suspended object when the suspended object was present and when it was absent, and normalizing this difference to the firing rate in the corresponding area near the standing object when this object was present.

Most of the object-induced fields in border cells had properties similar to those of object-vector fields with no border activity. The fields were confined and distributed across a wide spectrum of angles, not deviating significantly from a uniform distribution (Fig. 6c; Rayleigh test, Z = 1.1, n = 13, P = 0.35). Their orientation with respect to the object could not be predicted by the orientation of the border field (Fig. 6d; circular correlation r = −0.39, n = 14 object-vector fields from 12 cells, P = 0.13), and there was no significant correlation between the distance between border field and border and the distance between object-vector field and object (Fig. 6d; distance from border to centre of border field vs. distance from object centre to object-vector field centre: r = 0.04, n = 14, P = 0.89). There was a significant correlation between the difference in orientation to wall and object and the difference in distance (Fig. 6e; r = 0.56, n = 14, P = 0.04), i.e. cells that differed with respect to orientation also tended to differ with respect to distance. The distance between the object and the centre of the object-vector field in the 12 cells ranged from 9 to 39 cm (Fig. 6c; mean ± s.e.m.: 21.9 ± 2.6 cm), whereas the distance from the object to the field boundary ranged from 0 to 27 cm (9.6 ± 2.1 cm). These ranges contrasted with the distance from the wall to the centre of the border field, which ranged only from 6 to 14 cm (Fig. 6c; mean ± s.e.m, 8.8 ± 0.2 cm). Altogether, the analyses suggest that a small fraction of the object-vector cells overlapped with the border-cell population. In those cells, there was no consistent vectorial relationship from firing field to border and object.

Finally, we asked if cells with responses to both borders and objects could be differentiated with regard to how they respond to elevated objects. Five of the 12 border cells that also passed the object-vector cell criteria were tested with suspended objects in addition to standing objects (Fig. 6 f,g). We quantified, for these cells, the difference in response to standing and suspended objects using the object-vector field at the standing object to define a template area with a given distance and orientation from the object, and then applying the same vector-defined area with reference to the suspended object (Fig. 6f). Corresponding areas were next identified for the no-object trial. The response to standing and suspended objects was then taken as the difference in firing rate on no-object and object trials in the template area defined by the suspended object, normalized to the firing rate in the template area for the standing object on the object trial. For the 21 non-border object-vector cells tested with suspended objects, the score was always positive (Fig. 6h; mean ± s.e.m., 1.1 ± 0.17), suggesting that firing rates increased reliably in the template area near the suspended object, whereas for the 5 border cells, the score was only −0.24 ± 0.09 (non-border vs. border: U = 334, n1 = 21, n2 = 5, P = 7.2×10^−4^, Mann-Whitney U-test), indicating no increase in firing compared to no-object trials in the vicinity of suspended objects. These observations raise the possibility that object-vector cells and border cells respond differentially not only to the extension of object dimensions but also to their vertical location and the degree to which they constrain the animal’s path.

## Discussion

We have shown that a distinct population of MEC cells encodes direction and distance from discrete objects, independently of the identity or location of the object in the environment. These cells differ from the object-encoding cells of the LEC, which fire only at the object location and not in the space between the objects. Object-vector firing fields are expressed instantly when new objects are encountered in new environments. The vector representation is allocentric, i.e. the firing fields are fixed in room coordinates, irrespective of the animal’s direction of movement, distinguishing them from the egocentrically tuned goal-vector cells of the hippocampus^8,9^. Object-vector cells are part of a broader MEC representation, with which it remains aligned even when grid and head-direction cells undergo changes in spatial and directional tuning, such as after relocation from one room to another. Taken together, these results identify an object-centered entorhinal metric of navigational space tightly integrated with the more distantly anchored framework enabled by grid cells and head-direction cells.

Object-vector cells provide a possible cellular basis for spatial mapping where objects rather than surrounding geometry serve as the reference frame. Unlike laboratory environments, real worlds are cluttered with objects. When objects maintain stable positions relative to a goal, animals have been observed to use them as references for navigation^35–40^. Based on this finding, theoretical studies have proposed the existence, in cortical networks, of allocentric vectorial representations where trajectories are derived from differences between vectors from the animal to perceived objects and vectors between such objects and goals stored in memory^41^. The present study shows that vectors to perceived objects are represented in a dedicated neural population in the superficial layers of MEC. This neural population interweaves with the network of grid cells and head direction cells that encodes position for space in a more distal geometric framework^21,26,52^, independently of the location of the objects. The continued alignment of these networks across environments suggests that the networks are strongly interconnected. Although their mode of interaction remains to be determined, one possibility is that object-vector cells receive distally anchored positional and directional input from grid cells^53^ and head direction cells^41^, and that this metric is anchored to landmarks by association with visual or other sensory signals, or signals from object-informative cells in LEC^33,34^ or elsewhere^54,55^. The strong reciprocal connections of MEC and LEC^56–58^ might enable such interactions.

Previous studies have identified object-responsive cells across a wide network of brain regions that include both LEC^33,34^ and areas connected with the LEC, such as the perirhinal^54^ and anterior cingulate^55^ cortices. These ‘object cells’ differ from the object-vector cells of the MEC in that they respond only when the animal is at the object, not when it perceives it from a distance. The cells also lack directional tuning. Thus, they cannot alone account for navigation in the space between objects. More direct evidence for cells with object-vector properties comes from a study of CA1 cells in which place cells were found to intermix with a sparse population of cells that fired repeatedly at specific distances and directions from a subset of objects placed in the recording compartment^45^. Earlier work has similarly shown that firing fields of some CA1 cells follow the location of an array of landmarks when the array is moved in a test arena^59^, and more recent work has identified subsets of CA1 place cells that fire consistently at or before mobile visual-tactile landmarks on a linear treadmill, regardless of the location of the landmarks on the belt^60^. While allocentric directional relationships were either not investigated or not quantified in these studies, the cells do seem to have properties in common with the object-vector cells of the MEC. But there are also likely to be important differences. Hippocampal cells often responded only to a subset of the objects, unlike their entorhinal counterparts, which developed fields unconditionally at every single object the mouse encountered. Moreover, hippocampal cells fired at a single location relative to each object, while the entorhinal object-vector cells frequently expressed firing fields in more than one direction from it. Finally, during foraging in the box, the firing fields of the hippocampal cells in many cases developed over multiple trials^45^, distinguishing them from entorhinal object-vector cells, which emerged instantly, not requiring any extended experience with the environment or the object. The apparently slow appearance of object-vector representations in the hippocampal population points to instantly-appearing object-vector cells in MEC as a possible source for training hippocampal cells secondarily to form vectors between often-visited goals and stable landmarks for subsequent storage in memory^41^. Such a training process would be reminiscent of how odour-selective cells in the LEC can, over many trials, entrain odour-selective cells in the CA1 ^61^.

We have characterized a population of object-vector cells in MEC that responds to a wide variety of spatially confined, tower-like objects. The exact range of objects and object shapes that elicit vectorial representations in this cell population remains to be determined but our study shows that the majority of object-vector cells fail to respond to spatially extended objects in the distal background, such as the walls of the recording box, whereas instead they respond to discrete objects that stand out from the monotonous planar surfaces of the environment. Obstruction of the animal’s upcoming path was not a condition for object-vector cells to respond, as these cells, unlike border cells, fired vigorously also to suspended objects that the mice could not reach. The distinction between confined objects and walls was not absolute, however, because approximately one-third of the border cells with firing fields alongside the wall of the box had additional circular fields at specific distances and directions from more point-like objects on the arena floor. This proportion is larger than expected if object-vector cells and border cells were independent populations and raises the possibility that object-vector and border cells are activated by a continuum of shapes and sizes, from point-like bodies to lengthy surfaces. The fact that border cells with responses to discrete objects did not develop additional fields when the discrete objects were out of reach may imply that object-vector cells and border cells are categories of a broader class varying along multiple axes and dimensions. Such a breadth of response profiles would enable MEC networks to perform positional vector calculations across a wide variety of naturalistic environments, where discrete landmarks often are more common than the long walls or edges of test boxes in the laboratory.

Because object-vector cells define a range of distances and directions from restricted locations, they are reminiscent of ‘boundary-vector cells’ predicted by theoretical models of spatial coding^42–44^. In these models, each boundary-vector cell fires at a specific distance and direction from an extensive boundary in the local environment. Inputs from such boundary-vector cells were proposed by the models to give rise to place cells in the hippocampus. Cells with properties similar to boundary vector cells have been reported in the subiculum^46,47^ but most of these cells fire preferentially along the boundaries of the environment and not primarily out in open arena space^47^. Furthermore, cells with boundary-related activity in the subiculum do not project to place cells in the hippocampus. Instead, place cells receive abundant inputs from the superficial layers of MEC, where boundary-vector cells with fields offset from the arena walls are not common^22,23,48^. Our study shows that the MEC instead has a high density of object-vector cells, which define positions anywhere in the local space based on vectors from discrete objects, including vertical surfaces. This representation is versatile not only because it operates universally across environments and covers positions all across the environment, but also because it is independent of the presence of lengthy monotonous boundaries in the environment.

## Acknowledgments

We thank A.M. Amundsgård, K. Haugen, K. Jenssen, E. Kråkvik, and H. Waade for technical assistance. The work was supported by two Advanced Investigator Grants from the European Research Council (GRIDCODE’, Grant Agreement N°338865; ENSEMBLE’ – Grant Agreement N°268598), a NEVRONOR grant from the Research Council of Norway (grant no. 226003), the Centre of Excellence scheme and the National Infrastructure Scheme of the Research Council of Norway (Centre for Neural Computation, grant number 223262; NORBRAIN1, grant number 197467), the Louis Jeantet Prize, the Körber Prize, and the Kavli Foundation.

## Author Contributions

Ø.A.H., M.-B.M. and E.I.M. designed experiments and analytic approaches; Ø.A.H. and E.R.S. collected data; Ø.A.H. performed analyses; M.-B.M. and E.I.M. supervised the project; Ø.A.H. and E.I.M. wrote the paper with input from all authors.

## Author Information

Reprints and permissions information is available at www.nature.com/reprints.

## Methods summary

Methods, along with any additional Extended Data display items, are available in the online version of the paper; references unique to this section appear only in the online paper.

### Online Methods

#### Subjects

Data were obtained from 8 male wild type black 6 mice at the age of 4-11 months. All mice were kept on a 12 hr light/12 hr dark schedule in a humidity and temperature-controlled environment. The mice were housed in single animal cages after implantation. Testing occurred in the dark phase. The animals were not deprived of food or water. Experiments were performed in accordance with the Norwegian Animal Welfare Act and the European Convention for the Protection of Vertebrate Animals used for Experimental and Other Scientific Purposes.

#### Surgery and electrode implantation

The mice were anesthetized with 5% isoflurane (air flow: 1.2 l/min) in an induction chamber. Upon induction of anesthesia, they received subcutaneous injections of buprenorphine (Temgesic) and Meloxicam (Metacam). The mice were then fixed in a Kopf stereotaxic frame for implantation. Local anesthetic Bupivacaine (Marcaine) was injected subcutaneously before making the incision. During surgery, isoflurane was gradually reduced from 3% to 1% according to physiological condition. The depth of anesthesia was monitored by testing tail and pinch reflexes as well as breathing.

Anesthetized mice were implanted with a single bundle of 4 tetrodes attached to a microdrive fastened to the skull of the animal. The tetrodes were targeted to MEC at an angle of 3-4 degrees relative to the bregma/lambda horizontal reference plane, with the tips pointing in the posterior direction. The tetrodes were inserted 3.1-3.3 mm lateral to the midline and 0.3-0.4 mm anterior to the transverse sinus edge, with an initial depth of 800 μm. The implant was secured to the skull with histoacryl and dental cement. One screw was connected to the drive ground.

Tetrodes were constructed from four twisted 17 μm polyimide-coated platinum-iridium (90%–10%) wires (California Fine Wire, CA). The electrode tips were plated with platinum to reduce electrode impedances to between 120 and 220 kΩ at 1 kHz.

#### Behavioral procedures

The mice were trained to forage for cookie crumbs in an 80 cm × 80 cm square and a 90 cm diameter circular compartment, both enclosed by 50 cm high walls. Thick dark blue curtains surrounded the recording arena, except for a slit to one side (Fig. 1a). Before testing, the mouse rested outside the curtain on a flowerpot on a pedestal covered with towels. Testing was performed at low light levels to encourage exploration. Between trial sequences, the mat covering the floor of the recording box was cleaned.

A typical trial sequence started with a trial where no object was present in the arena, followed by one in which a tower-shaped object was placed at a semi-randomly varied location, with a bias towards the box centre on the first trial (to capture fields with large offsets from the object). The object used for standard trials (Fig. 1 and Fig. 2) was usually Object 7 in Extended Data Fig. 2, a 5-cm wide, 20-cm tall cylinder-shaped object. For subsequent experiments, objects were selected from a pool of tower-like modified rectangular prisms or cylinders ranging in size between 3 and 7 cm in width and 9 and 20 cm in height for the prisms and between 3 and 8 cm in diameter and 20 and 35 cm in height for the cylinders. All of these objects were substantially taller than the mouse. In addition, on selected trials we used a flattened cylinder (11 cm in diameter, height of 0.5 cm; Object 13 in Extended Data Fig. 2), or a block that had the shape of a wall (Object 6 in Extended Data Fig. 2; 50 cm long, 0.5 cm wide, 50 cm high).

In order to verify that any observed change in neural activity between the first and the second trial was tied to the location of the object, the object was displaced in a pseudo-random fashion on a third trial in the sequence (only in those experiments where object-responsive cells were present on the preceding trial). In a few cases, not only the location but also the identity of the object was changed on the third trial (see below for tests with different object identities). The average object displacement on the third trial was 19.4 ± 1.1 cm (mean ± s.e.m.). Each trial lasted approximately 15 minutes. Trials were typically spaced by a few minutes, during which the experimenter clustered and inspected recorded cells and placed the object in a new location. The mouse was in most circumstances not removed from the arena between the trials.

In most experiments, only one object was placed in the arena, although tests with multiple simultaneously presented objects (2-6, mostly 2 or 3) were also conducted. In a subset of tests, cells were recorded in two rooms – one familiar and one new to the mouse. A few cells were recorded in both rooms also after the initial test in the second room. Only novel objects were presented in the novel room. Both novel and familiar objects were chosen from the pool of objects shown in Extended Data Fig. 2. Tests in familiar and novel rooms were consecutive, with an interval of 10-15 min for transport and preparation. During preparation for recording, the mouse rested in a flowerpot on a pedestal outside the curtains.

In a subset of the experiments, the mouse was tested with a suspended object, out of reach to the animal (Object 9 in Extended Data Fig. 2). On these tests, a blue plastic tube with a diameter of 7 cm was taped to the wall, with the lower end approximately 15 cm above the floor level. In each individual experiment, the mouse was observed to make sure that the elevation was sufficient to not obstruct the animal’s movement in any way. The suspended object was presented at the same time as a standing object elsewhere in the arena.

Finally, while the majority of experiments were performed with lights on, objects were also presented in darkness in a few instances. The mice first explored the arena with an object present and curtains fully enclosing the arena. Subsequently, the lights were turned off and the mouse explored the arena and the object in darkness. The mouse was not taken out of the arena between the light and dark trials. Each trial lasted approximately 15 minutes.

#### Recording procedure

Data collection started 1-2 weeks after implantation of the tetrodes. During recording, the animal was connected to an Axona data acquisition system (Axona Ltd., Herts, U.K.) via an AC-coupled unity-gain operational amplifier close to the animal’s head, using a light-weight counterbalanced multiwire cable from both implants to an amplifier. Unit activity was amplified 3000-14,000 times and band-pass filtered between 0.8 and 6.7 kHz. Triggered spikes were stored to disk at 48 kHz with a 32-bit time stamp. An overhead camera recorded the position of two light-emitting diodes (LEDs) on the head stage, each at a sampling rate of 50 Hz. The diodes were separated by 3 cm and aligned with the body axis of the mouse. To sample activity at multiple dorso-ventral MEC positions, the tetrodes were lowered in steps of 25-50 μm between trial sequences after all relevant tests had been completed. Recordings started as soon as the tetrodes were judged to be in MEC, using theta modulation and presence of spatially or directionally modulated cells as criteria in addition to tetrode depth.

#### Spike sorting and cell classification

Spike sorting was performed offline using graphical cluster-cutting software (tint; Neil Burgess and Axona Ltd.). Spikes were clustered manually in two-dimensional projections of the multidimensional parameter space (consisting of waveform amplitudes), using autocorrelation and cross-correlation functions as additional separation tools and separation criteria. Cluster separation was determined by calculating distances between spikes of different cells in Mahalonobis space (Extended Data Fig. 5). Noise in the vicinity of clusters was expressed as the L ratio (Extended Data Fig. 5). Clusters on successive recording trials were identified as the same unit if the locations of the spike clusters in cluster space were stable.

#### Firing-rate maps and head-direction tuning curves

Position estimates were convolved with a 35-point Gaussian window and *x*,*y*-coordinates were sorted into 2 cm × 2 cm bins. Spike timestamps were matched with position timestamps. Only spikes collected at instantaneous running speeds above 3 cm/s were included. Firing rate distributions were determined by counting the number of spikes and assessing time spent in 2 cm × 2 cm bins of the firing rate maps and in directional bins of 5 degrees in tuning curves for head direction. The distributions were smoothed with a two-dimensional Gaussian Kernel with standard deviation of 2.5 bins (5 cm) in both the *x* and the *y* direction for the rate maps and with a Gaussian filter with a standard deviation of 2 bins (10 degrees) for the head-direction maps.

#### Definition of object-vector cells

We detected firing fields in the rate map by iteratively applying the Matlab “contour” function (Mathworks.inc), starting from the cell’s peak firing rate until reaching two times the standard deviation of the firing rates of all bins in the rate map. Firing fields were defined as contiguous areas within a contour of at least 16 bins and with a peak firing rate of at least 2 Hz. Cells that after insertion of the object expressed firing fields that were not present on the no-object trial, were selected for further analysis. We distinguished between cells that fired at the object (object cells; 0-4 cm from the centre of the object) and cells whose firing fields were offset by more than 4 cm from the object. The focus of the analyses was on the latter category, which accounted for nearly all object-responsive cells.

For cells with fields that were offset by more than 4 cm from the object centre, the vector-relationship between the cell’s spatial firing and the object was expressed in a “vector-map” in which firing rate was measured as a function of distance and direction from the object location. With the centre of the object as the reference, the arena was divided into directional bins of 5 degrees’ width, and for each directional bin, the time-normalized firing rate was measured in distance bins of 2 cm (Fig. 1c). The resulting distance-by-direction matrix was smoothed with a two-dimensional Gaussian kernel using a standard deviation of 2 bins (10 degrees, 4 cm). In the vector map, 0 degrees was defined as east of the object in the room frame (Fig. 1 a,c). The vector map on the object trial was then compared with that of subsequent trial where the object was moved to a new location. Object-vector cells were defined as cells that had a correlation between vector maps on object and displaced-object trials that exceeded chance levels determined by repeated shuffling of the experimental data (see below). In addition, for a cell to be accepted as an object-vector cell, the vector-map correlation between object and displaced-object trials was required to be higher than between the no-object trial and either of the object trials, spatial information on the object trial had to exceed the 95^th^ percentile of the shuffled data, and the peak firing rate had to be at least 2 Hz. Vector maps for the no-object trial were made using the respective object locations on the object trials as references.

Shuffling was conducted with 100 permutations for each of the object trials. For each permutation, the entire sequence of spikes fired by the cell was time-shifted along the animal’s path by a random interval between on one side 20 s and on the other side 20 s less than the length of the session, with the end of the session wrapped to the beginning. Time shifts varied randomly between permutations and between cells. Vector maps were constructed for each permutation, with the object location of the experimental data as the reference in all permutations. Surrogate vector maps from the object trial were next correlated with surrogate vector maps from the displaced-object trial. The 99^th^ percentile correlation of the distribution of all vector-map correlations from all 120 cells was taken to be the chance level.

Vector maps were also used to determine the similarity of responses to different objects in experiments where multiple objects were present in the arena. For each cell, a vector-map was constructed around each object and fields were detected in the vector maps in the same way as for the firing rate maps. In these analyses, each vector map would contain multiple fields – at least one per object. We compared vector maps for different pairs of objects and defined the difference in orientation and distance of vectors to the two objects as the difference in orientation and distance of the two fields in the pair of maps that had the nearest peak-to-peak distance.

#### Egocentric directional tuning

Egocentric directional tuning curves were constructed according to the procedure described in Sarel et al.^9^. Briefly, tuning curves were constructed with movement (heading) direction bins of 20 degrees relative to the object location, and with zero degrees defined as moving toward the object and ±180 degrees moving away from the object. Curves were smoothed over 1.5 bins (30 degrees) with a Gaussian filter. We defined the egocentric directionality index as:

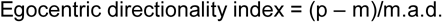

where p is the peak of the curve, m is the median of the curve, and m.a.d. is the median absolute deviation. Again we defined a 99^th^ percentile threshold by time-shifting the spikes 100 times for each cell and subsequently calculating egocentric directionality for each of these data.

#### Definition of grid cells

The spatial periodicity of each rate map – the cell’s grid score – was determined by calculating a spatial autocorrelogram^26^. For each cell, a grid score was determined by taking a central circular sample of the autocorrelogram, with the central peak excluded (the central peak was defined as 100 or more contiguous pixels of 1.5 × 1.5 cm^2^ above a fixed threshold of r > 0.1), and comparing rotated versions of this sample^26,62^. The Pearson correlation of the circular sample with its rotation in *α* degrees was obtained separately for angles of 60 and 120 on one hand and 30, 90 and 150 on the other. The cell’s grid score was defined as the minimum difference between any of the elements in the first group (60 and 120 degrees) and any of the elements in the second.

A cell was defined as a grid cell if its grid score exceeded a chance level determined by repeated shuffling of the experimental data (100 permutations per cell). For each permutation, the entire sequence of spikes fired by the cell was time-shifted along the animal’s path by a random interval between on one side 20 s and on the other side 20 s less than the length of the session, with the end of the session wrapped to the beginning. Time shifts varied randomly between permutations and between cells. If the grid score from the recorded data was larger than the 99^th^ percentile of grid scores in the distribution of shuffled data from all cells, the cell was defined as a grid cell. As an additional criterion we required the cell to have a peak firing rate of at least 2 Hz.

#### Template grid patterns from object-vector cells with multiple fields

For cells with two or more object-vector fields, we constructed a regular grid lattice extrapolated from the positions of the two object-vector fields. Template fields were modelled as circular areas centred at vertices in the grid lattice, and with size equal to the mean area of the two object-vector fields. A Z-score was calculated by first determining the difference between the mean firing rate inside the extrapolated template areas and the mean firing rate outside all projected and real firing fields, and then dividing this difference by the standard deviation of the firing rate of all bins in the rate map.

#### Analysis of head direction cells

The animal’s head direction was determined for each tracked sample by plotting the relative positions of the two LEDs onto the horizontal plane. The directional tuning function for each cell was obtained by plotting the firing rate as a function of the animal’s heading direction, divided into bins of 5 degrees and smoothed with a Gaussian moving average of 2 bins on each side. Directional tuning was estimated by computing the length of the mean resultant vector (mean vector length) for the circular distribution of firing rates. For a cell to be included as a head direction cell, its mean vector length needed to pass the 99^th^ percentile threshold of the mean vector length in the shuffled version of the same data. Shuffling was performed by shifting spike times at random intervals along the path in the same way as for grid cells.

#### Analysis of border cells

Border cells were identified by computing a border score for each cell^23^. Firing fields were defined as areas corresponding to a minimum of 9 neighboring bins (1 bin = 2 cm) of the smoothed rate map where firing rates were higher than 20% of the cell’s peak firing rate. The cell’s border score was expressed as the difference between the maximal length of a wall touching on any single firing field of the cell and the average distance of the field from the nearest wall, divided by the sum of those values. Border scores ranged from - 1 for cells with infinitely small central fields to +1 for cells with infinitely narrow fields that lined up perfectly along the entire wall. For circular arenas, the rate map was first transformed into a square image using a polar transform with the centre of the box as the reference. The resulting polar image represented distance from the wall (in cm) on one axis and distance along the perimeter or the wall (in angles) on the other. The border score was then computed as for square arenas. Since border cells often cover half of the arena in a circular environment, a border score of +1 would be assigned to cells with infinitely narrow fields lining up along the wall of half the box, or covering 180 degrees in the polar-image. After border scores were computed for each cell, border cells were identified as cells in which the border score exceeded the 99^th^ percentile of the distribution of border scores in shuffled versions of the same data. Shuffling was performed by shifting spike times at randomly chosen intervals along the path in the same way as for grid cells. In addition, for a cell to be defined as a border cell, we required the cell to have a peak rate of at least 2 Hz and a spatial information content that exceeded the 95th percentile for spatial information on the no-object trial.

#### Spatial information content and spatial coherence

Information content was determined for each rate map by computing the spatial information rate as

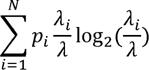

where λ_i_ the mean firing rate in the *i*-th bin, λ the overall mean firing rate and *p_i_* the probability of the animal being in the *i*-th bin (occupancy in the *i*-th bin/total recording time). Spatial information in bits/spike was obtained by dividing the information rate with the mean firing rate of the cell. Spatial coherence was calculated for each cell as the z-transform of the correlation between the firing rates of each bin and the averaged firing rates of the 8 nearest neighbours of that bin.

#### Coherence between directional or spatial rate distributions

We quantified the degree to which orientation of different cell types shifted coherently between trials in different rooms by performing circular cross-correlations on their directional or spatial rate distributions. For each head-direction cell, we cross-correlated the directional tuning curves obtained from the two recording rooms, using a bin size of 5 degrees. For object-vector cells, we cross-correlated the object-centered allocentric directional tuning curves - where firing rate was expressed as a function of orientation relative to object - across trials in the same two rooms, also using a bin size of 5 degrees. For grid cells, we rotated the rate map in the first room in steps of 5 degrees and correlated this map with the rate map of the other room for each step. For all cell types, we identified the shift between the two pairs of distributions or maps that gave the peak correlation. Two of the object-vector cells with two firing fields had symmetrically arranged fields, one on each side of the object, yielding two clear peaks in the cross-correlation. In these two cases we determined the shift of the maps by taking advantage of consistent differences in field sizes and distances between the two fields and the object. For each pair of concurrently recorded cells, we noted the difference in the angles that resulted in peak correlation. We compared the distribution of these differences with the distribution of pairwise differences obtained by randomly shifting the tuning curves in a circular manner while maintaining the shapes of the tuning curves.

#### Response to suspended object

Border cell and object-vector cell response to suspended objects was compared by first identifying firing fields in the vicinity of a standing object, as described above. The field area and vector from object to field-center was then transferred to the projection of the suspended object on the floor surface to act as a template for an expected field (Fig. 6f). Mean firing rates were measured in both of these areas in the presence of the objects and on no-object trials. This procedure was also used to compare how well the vector between a firing field of an object-vector cell and the object in one object trial predicted the position of a field in a trial where the object was displaced (Extended Data Fig. 3a). We defined a suspended-object response score as:

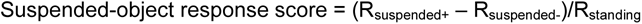

where R_suspended+_ is the mean firing rate in the template area associated with the suspended object, R_suspended−_ is the mean firing rate in the template area when the suspended object is absent, and R_standing_ is the mean firing rate of the original field in the vicinity of the standing object.

#### Histology and Reconstruction of Recording Positions

The tetrodes were not moved after the final recording session. The mouse received an overdose of Pentobarbital and was perfused intracardially with 9% saline and 4% formaldehyde. The brain was extracted and stored in 4% formaldehyde. Frozen, 30mm sagittal sections were cut, mounted on glass, and stained with cresyl violet (Nissl). The final position of the tip of each tetrode was identified on photomicrographs obtained with Axio Scan.Z1 microscope and Axio Vision software (Carl Zeiss, Germany) (Extended data Fig. 1).

#### Statistical tests

All statistical tests were two-sided. We used Mann-Whitney U tests for independent group comparisons and Wilcoxon signed rank tests for paired tests. Correlations were determined using Pearson’s product-moment correlation coefficients. P-values for Pearson’s correlations were computed using a Student’s *t* distribution for a transformation of the correlation (Matlabs ‘corr’ function). Binomial tests were used to determine expected probabilities of observing reported counts. No statistical methods were used to pre-determine sample sizes but our sample sizes are similar to those reported in previous publications.

---

**Extended Data Figure 1.**
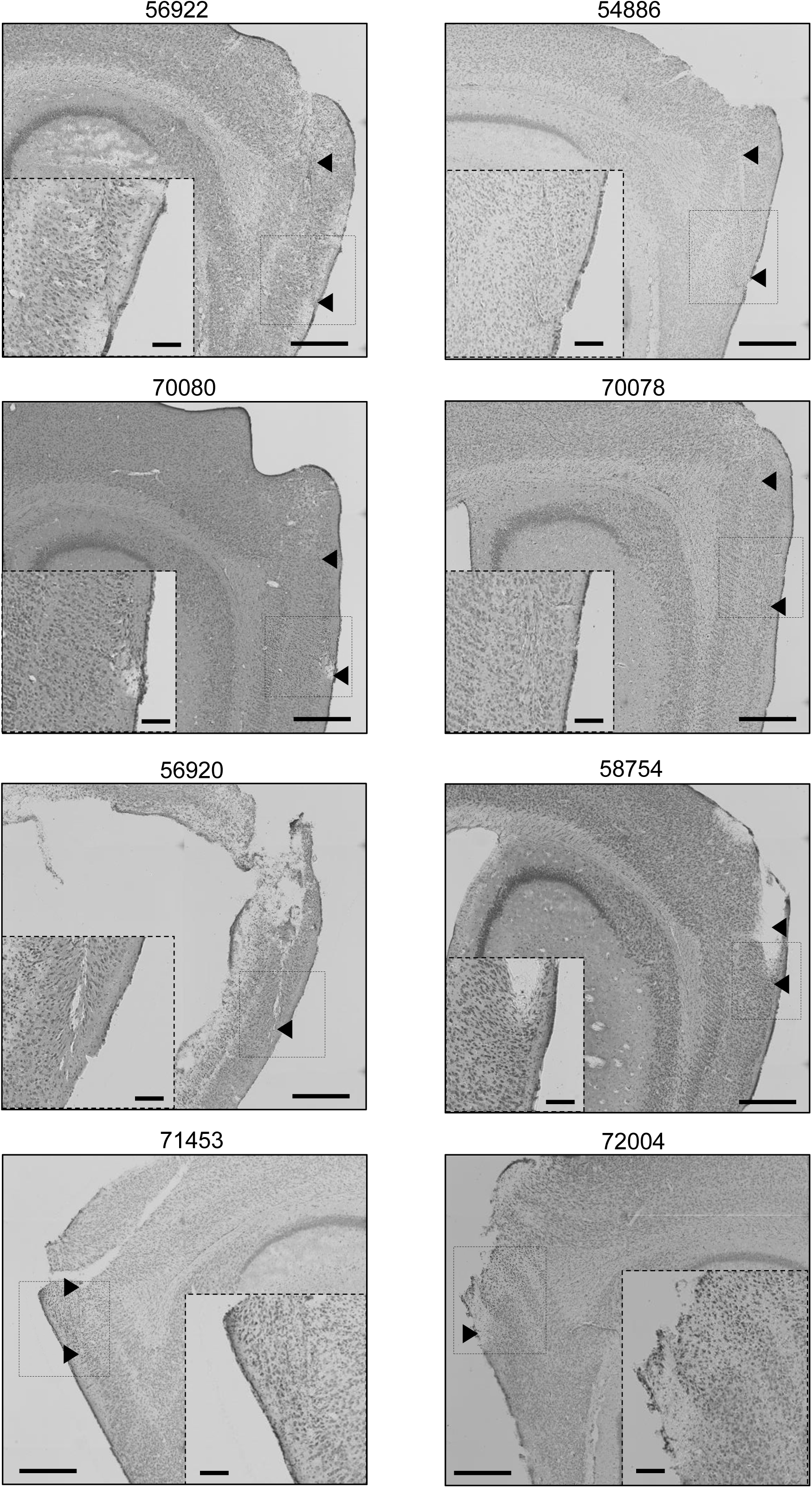
Recording locations in MEC. Nissl-stained sagittal brain sections showing tetrode locations for the 8 mice used for experiments. Pairs of black arrows indicate dorsoventral range of recording locations. Insets show the end of the tetrode track at higher spatial resolution (magnified area is indicated in the low-resolution images). Small scale bars: 250 µm, large scale bars: 500 µm. The hippocampal area of mouse 56920 was damaged during extraction of the brain but the tetrode tracks can still be identified in the superficial layers of the MEC also in this animal.

**Extended Data Figure 2.**
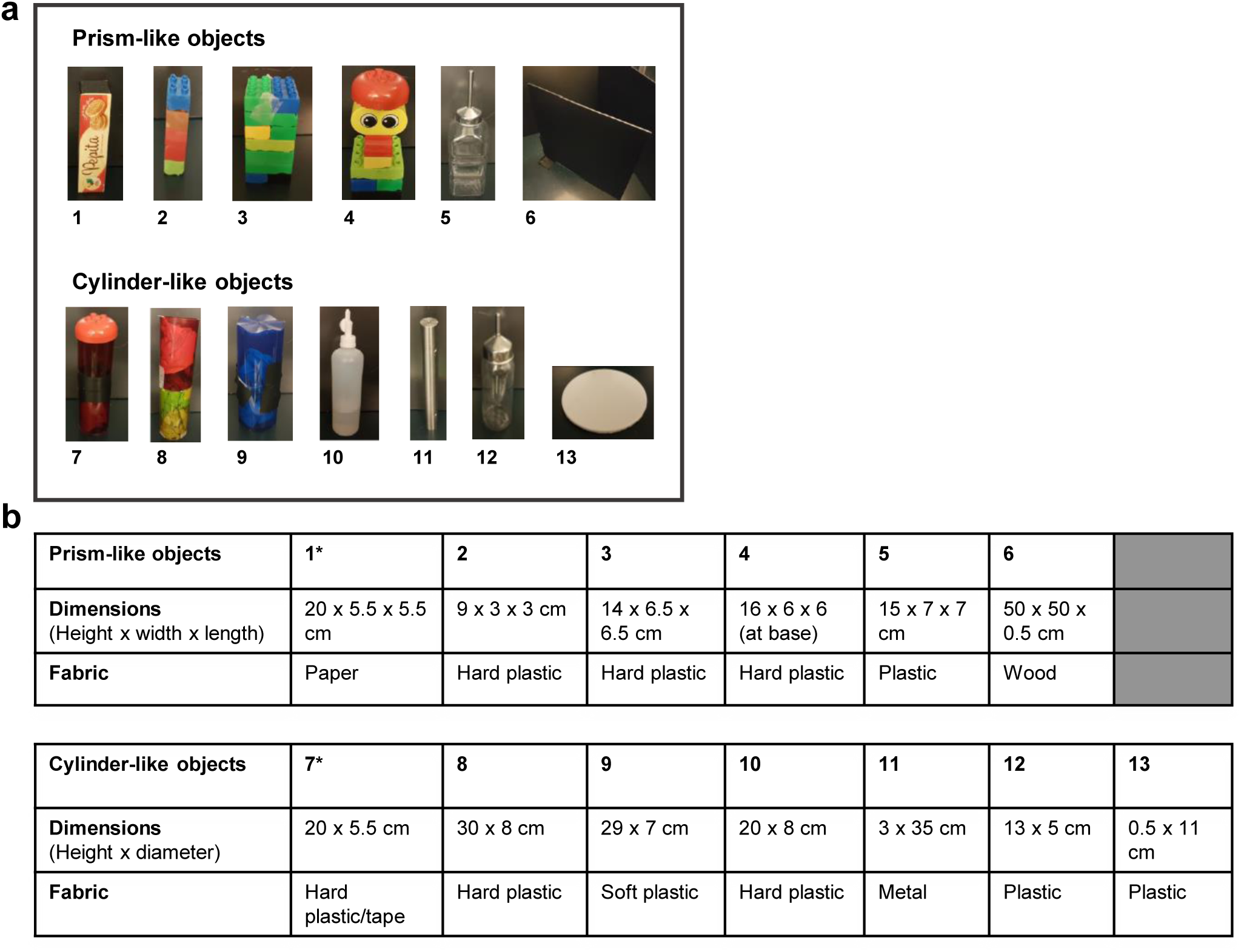
List of all objects used to identify object-vector cells. **a)** Photographs of all objects used in experiments with object-vector cells. Objects had either a prism or cylinder-like appearance, with some modifications, and varied in height and width. Object 6 – a wall insert – deviated from the tower-like shape of the other prisms. Object 13 was almost flat and deviated from all other objects, which were taller than the mouse. **b)** Table showing shape, dimensions, colour and fabric of each object in a. Objects are numbered as in a.

**Extended Data Figure 3.**
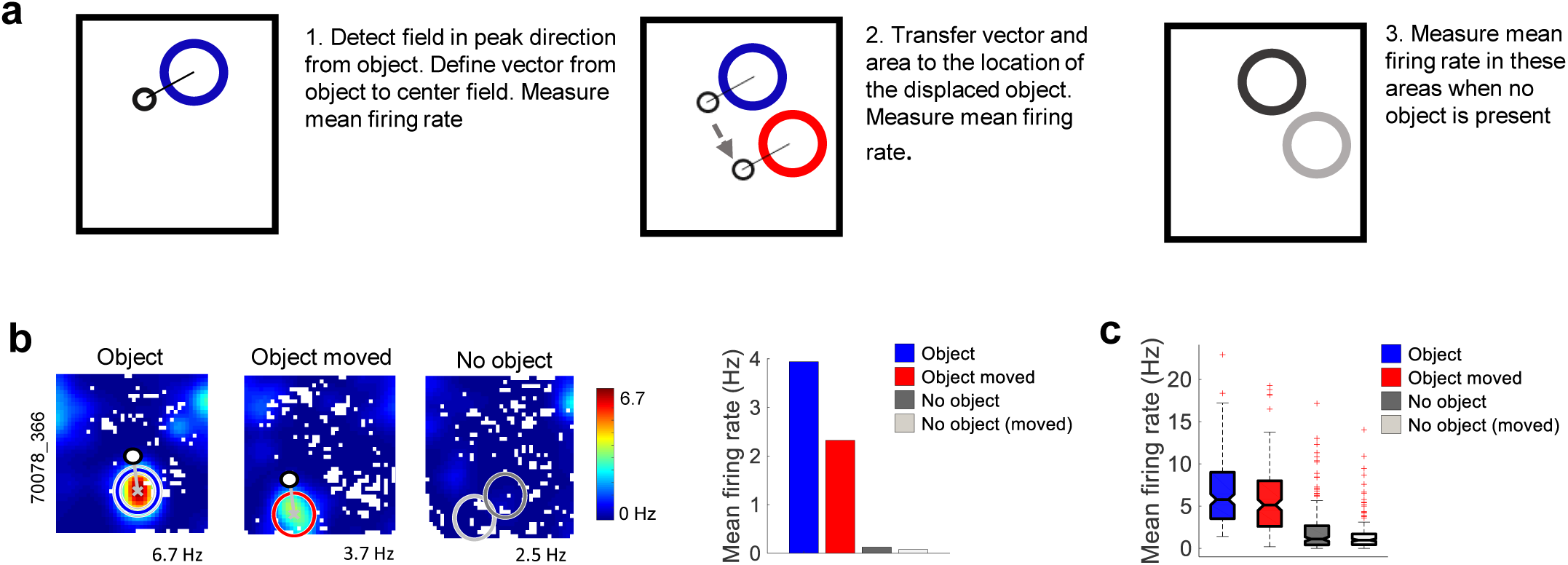
Object-vector cells have fixed distance and direction relationships to objects. **a)** Procedure for comparing object-vector fields across object locations. The area of an object-vector field and the vector between the centre of the field and the centre of the object were transferred as templates to the position of the displaced object. Mean firing rates were measured in both the original and the template area when objects were present and when no object was in the enclosure. **b)** Application of this procedure on an example object-vector cell. Mean firing rates within original and template areas are shown in colour-coded rate maps and as a histogram for trials with no object, object, and displaced object. Peak rates (horizontal) and mouse and cell number (vertical) are indicated on the rate maps. **c)** Box plots showing, for all object-vector cells, the distribution on object trials of mean firing rates in regions with similar distance and direction relationships to the object, as well as firing rates in the same areas when an object was not present. On trials with displaced objects, object-vector cells showed reliable increases in firing rate in the template area defined by the cell’s vector relationship to the object on the original object trial.

**Extended Data Figure 4.**
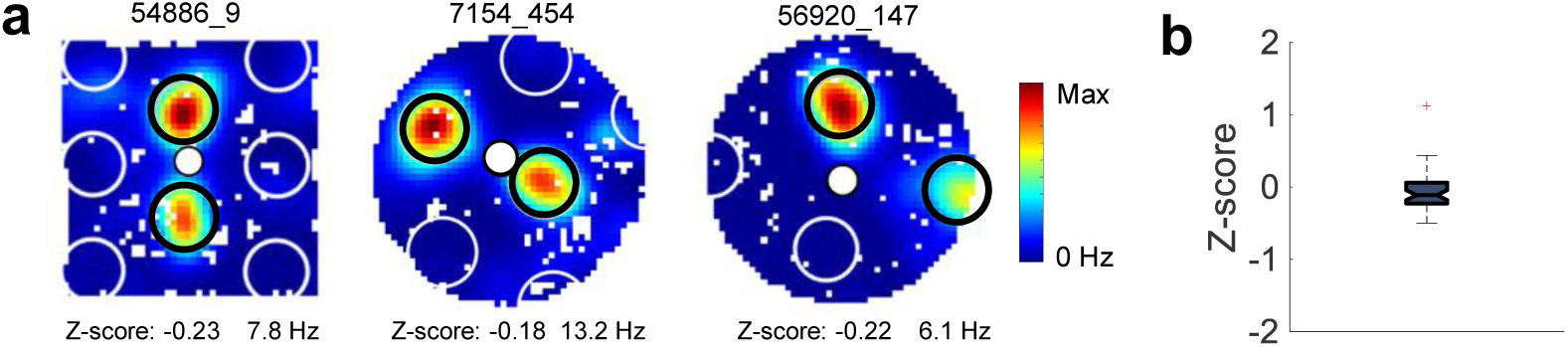
Object-vector cells with multiple fields are not grid cells. **a)** Colour coded rate maps of three object-vector cells with two object-vector fields. Object-vector fields are indicated by black open circles. Small filled white circles represent objects. Large white and open circles indicate template areas in a regular grid lattice extrapolated from the positions of the two object-vector fields. A Z-score is calculated by first determining the difference between the mean firing rate inside the extrapolated areas (large white and open circles) and the mean firing rate outside all projected and real firing fields, and then dividing this difference by the standard deviation of the firing rate. Z-scores, as well as peak firing rates, are indicated below each of the three example maps. Mouse and cell number are indicated at the top. **b)** Box plot showing Z-scores, calculated as in a, for the entire population of object-vector cells with multiple object-vector fields. Mean scores (± s.e.m.) were low (−0.06 ± 0.04). Fluctuation of Z-scores around 0 suggests that the two (or three) fields of the object-vector cells are not part of a regular grid pattern.

**Extended Data Figure 5.**
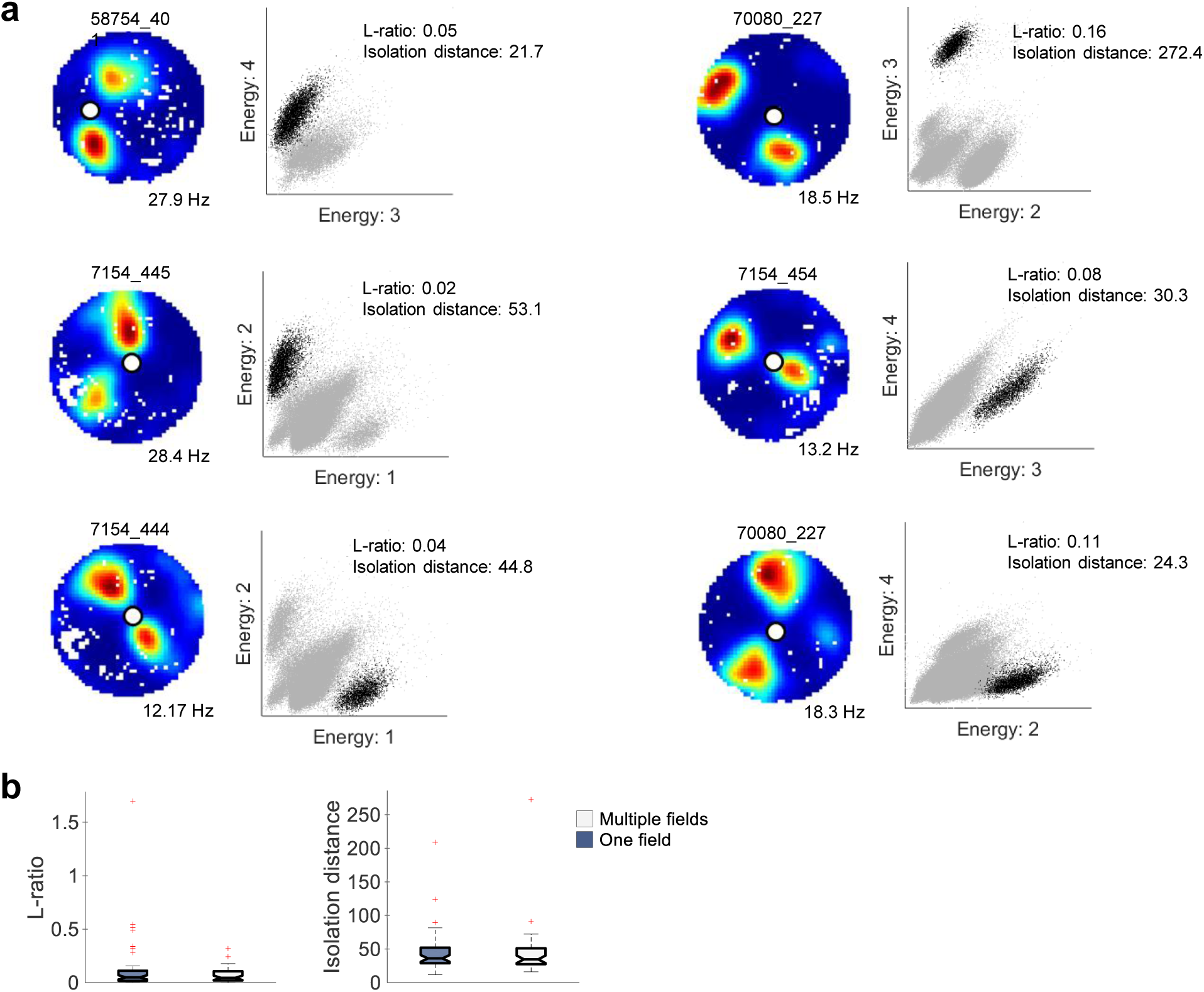
Spike clusters of object-vector cells with two discrete firing fields. Separation of spikes from object-vector cells with two firing fields from spikes of other simultaneously recorded MEC cells. **a)** Examples of cell separation in two-dimensional projections of multidimensional cluster diagrams. First and third columns: Color-coded rate maps showing two distinct object-vector cells, with mouse and cell numbers (top) and peak rates (bottom) indicated. Second and fourth columns: Scatter plots showing relationship between energy (square of signal) for spikes recorded from two selected electrodes of a tetrode in the recording containing the object-vector cell in the rate map to the left. Pair of electrodes is indicated on axis labels. Each dot represents one sampled signal. Clusters are likely to correspond to spikes originating from the same cell. The cluster giving rise to the rate map to the left of each scatterplot is shown in black; remaining signals in grey. L-ratio and isolation distance^63^ for cluster in black are indicated above the scatter plot. Note clear separation of the object-vector cell from other spikes, suggesting it is unlikely that second fields reflected contamination of spikes from other cells. **b)** Box plots showing distribution of L-ratio and isolation distance for all object-vector cells with one field (dark blue boxes), and all with two or three fields (light grey boxes).

**Extended data. Figure 6.**
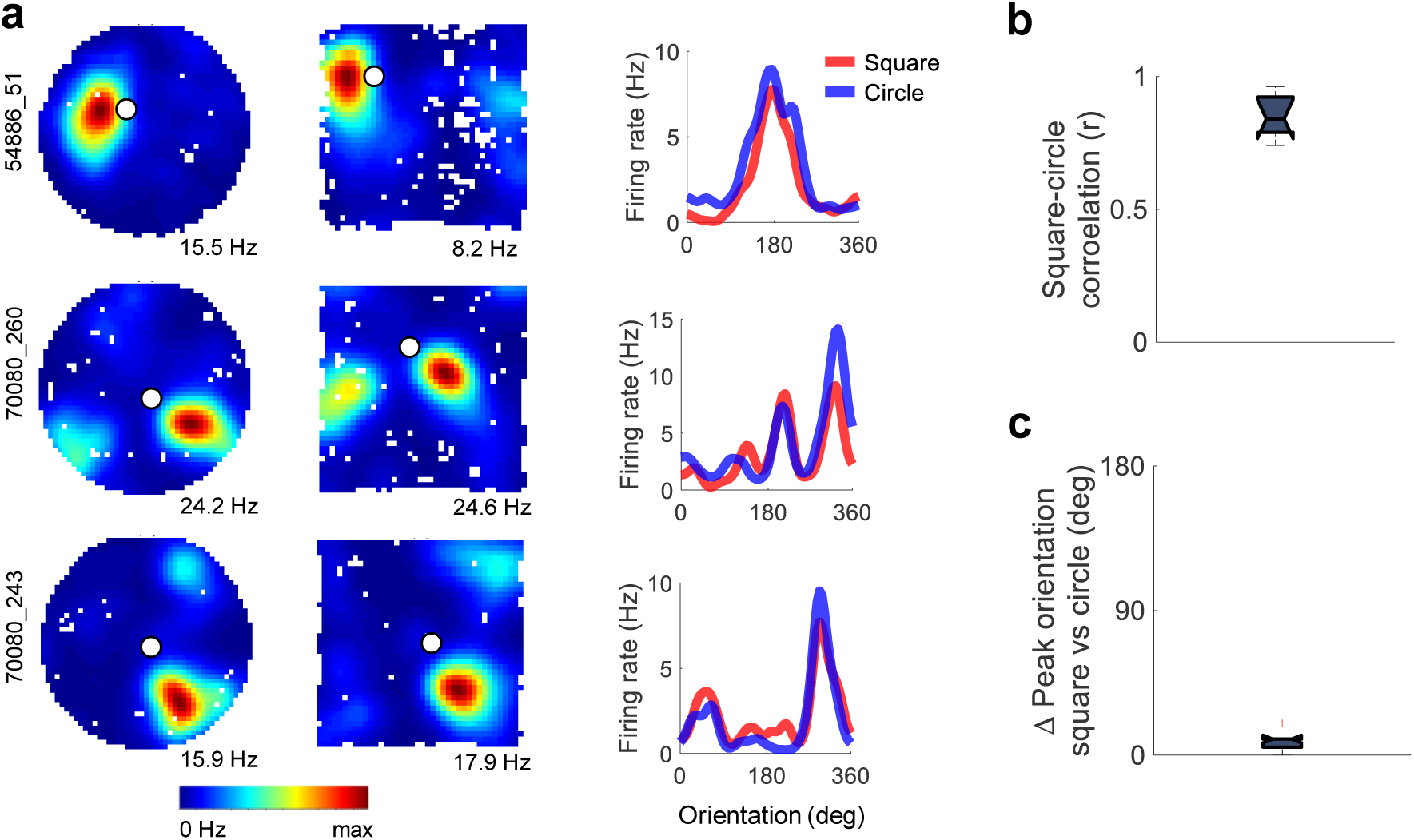
Orientation of object-vector fields is independent of the geometry of the environment. Experiment where 8 cells from 3 animals were recorded successively in a circular and a square recording box. The boxes were placed at the same location in the recording room and cues external to the box were identical. **a)** Left: colour-coded rate maps of 3 cells recorded first in the circle, then in the square. Peak rates are indicated below the rate maps, mouse and cell number to the left. Middle: directional tuning curves for the same 3 cells, with firing rates shown as a function of allocentric direction relative to the object. Tuning curves were smoothed with a Gaussian kernel with a standard deviation of 2 bins (10 degrees). **b)** Box plot showing correlation between orientation tuning curves in square and circular environments for all 8 cells. **c)** Box plot showing difference in peak direction of orientation tuning curves between square and circle for all 8 cells.

**Extended Data Figure 7.**
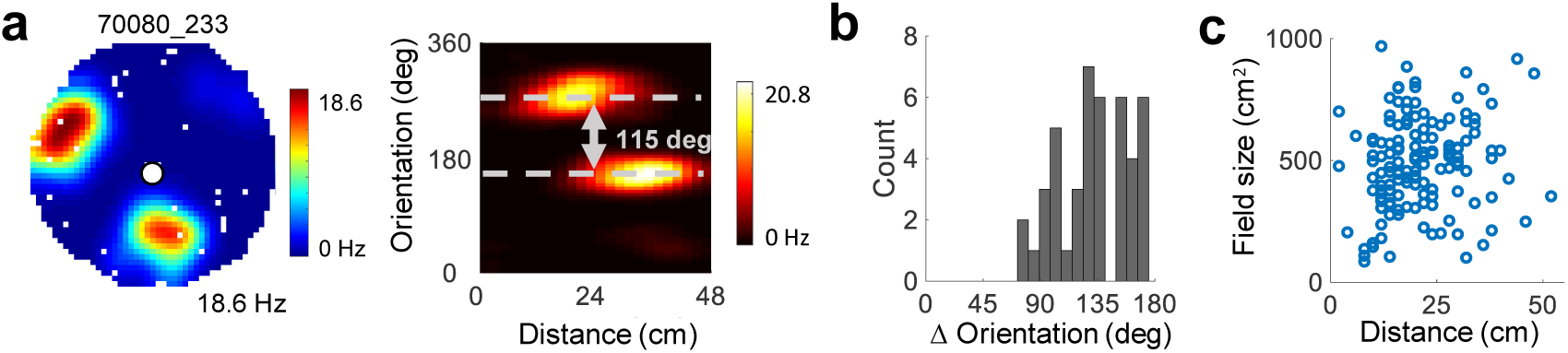
Orientation and size of object-vector fields for cells with one or more fields. **a)** Left: Colour-coded firing rate map of cell with two firing fields. White circle indicates location of object. Peak rate and mouse and cell number are indicated. Right: Colour-coded vector map showing firing rate as a function of distance and orientation from the object for the same cell as in the rate map. Radial difference between the two peaks is indicated. **b)** Distribution of orientation difference between pairs of fields for all cells with multiple object-vector fields. **c)** Field size plotted against distance from the object to the peak of the object-vector field (all object-vector cells).

**Extended Data Figure 8.**
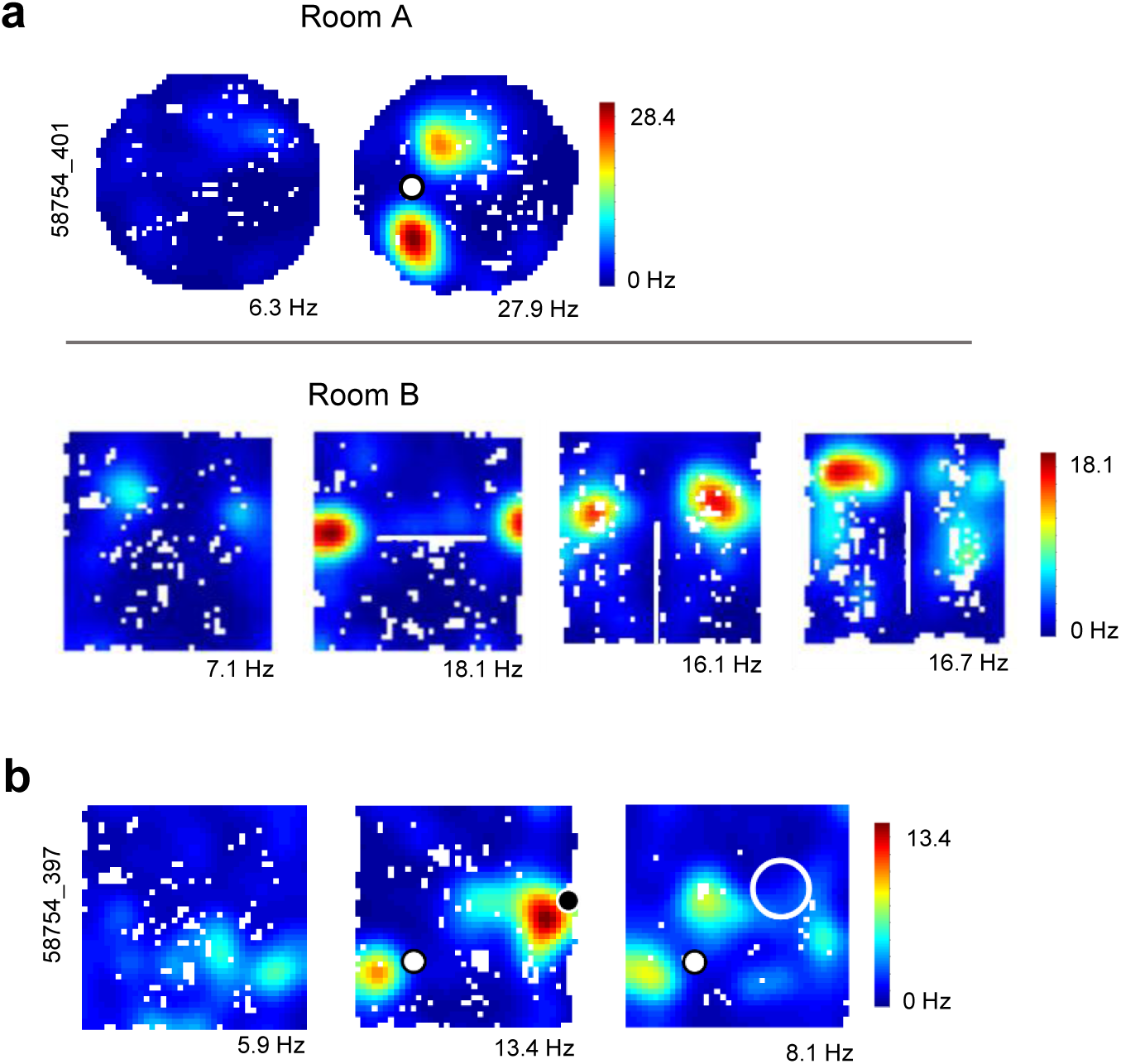
Object-vector response to flattened objects. **a)** Top row: colour-coded rate maps for object-vector cell recorded with no object and with an object (white circle; Object 13 in Extended Data Figure 2; room A). Peak rates (horizontal) and mouse and cell number (vertical) are indicated. The cell has two discrete firing fields when the object is present. Bottom row: same cell recorded in a different box in a different room (room B), with either no object (leftmost) or with a wall insert (right 3 panels; Object 6 in Extended Data Figure 2). The wall (white stripes in the rate maps) was placed centrally in the box with passage at both ends, or in touch with the wall of the box, with passage at only one end. **b)** Cell recorded with no object (left), with both a standing and a suspended object (middle), or with a standard and a flat object (flat object is Object 13 in Extended Data Fig. 2). Filled white circle indicates standing object, filled black circle suspended object, and open white circle flat object. Firing fields were detected to the left of each object.

**Extended data Figure 9.**
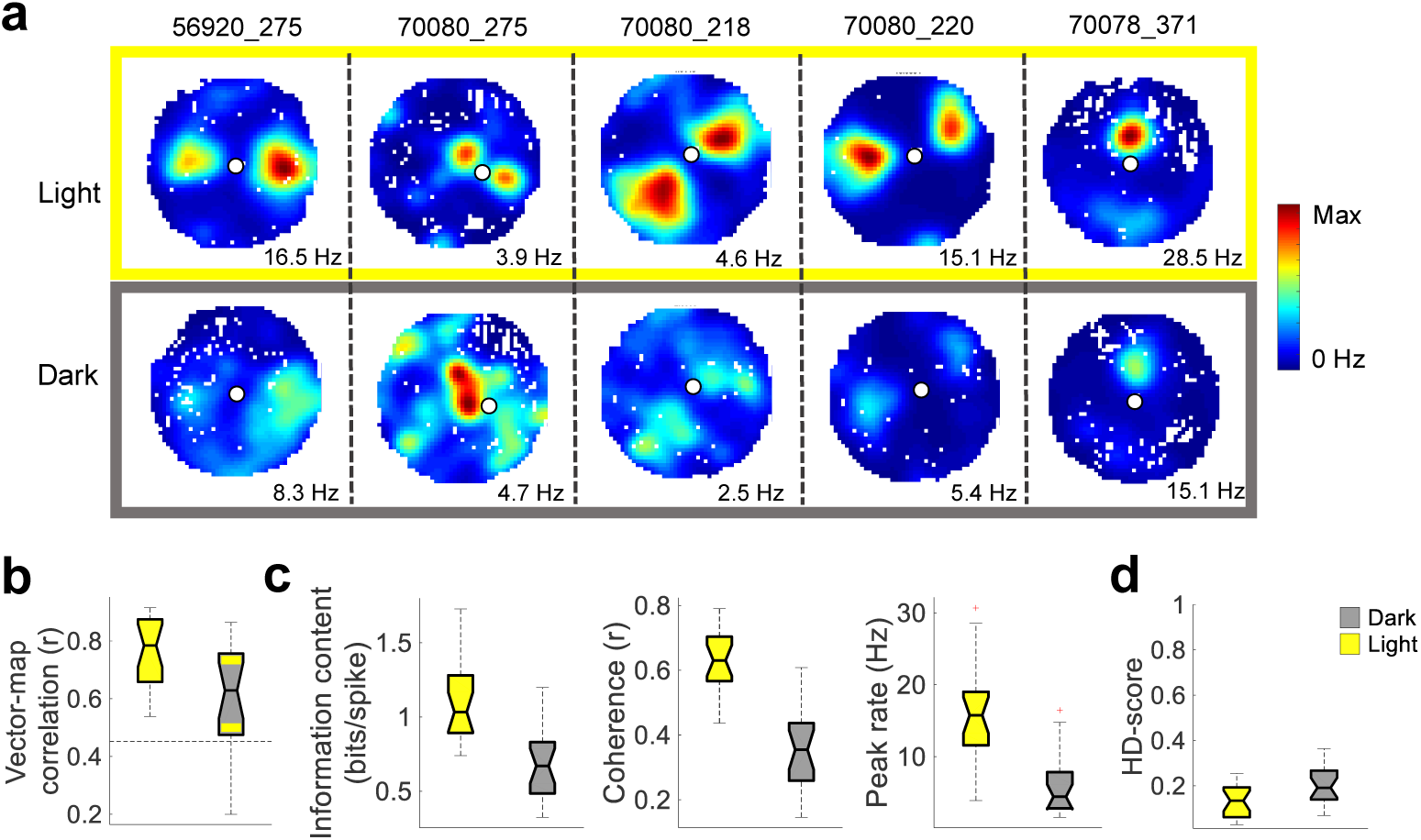
Object-vector cells lose accuracy in darkness. **a)** Colour-coded firing rate maps from 5 example cells recorded successively in light and complete darkness (top and bottom rows, respectively). Mouse and cell number are shown at the top of each column, peak rates below each rate map. Colour scale is similar for each column of rate maps. **b)** Box plot showing distribution of vector-map correlations across pairs of trials recorded either successively in light (yellow) or first in light and then in darkness (grey). Stippled line marks the 99^th^ percentile correlation threshold. **c)** Box plots showing distribution of spatial information content, spatial coherence, peak firing rate and mean firing rate in light and in darkness. All four measures decreased from light to darkness (spatial information content: Wilcoxon signed rank test: W = 227, n = 21, P = 1.1×10^−4^, spatial coherence: W = 231, n = 21, P = 6.0×10^−5^, peak firing rate: W = 230, n = 21, P = 6.9×10^−5^, and mean firing rate: W = 212, n1 = n2 = 95, P = 8.0×10^−4^). **d)** Box plot showing distribution of head-direction (HD) scores in light and in darkness. Head direction tuning increased significantly in the absence of visual cues (Wilcoxon signed rank test: W = 36, n = 21, P = 0.005).

**Extended Data Figure 10.**
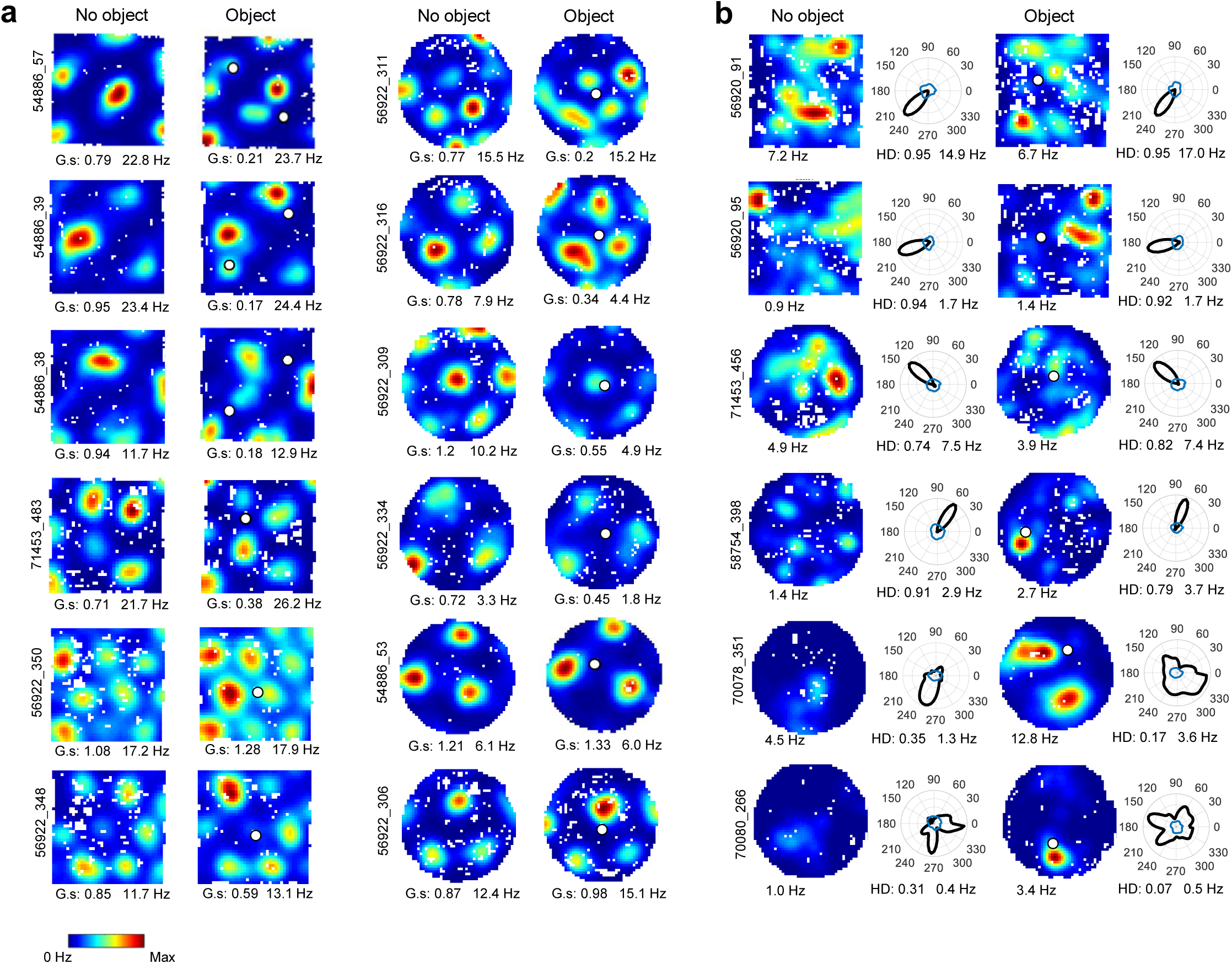
Effects of objects on grid cells and head direction cells. **a)** Colour-coded rate maps showing 12 representative grid cells recorded in the absence (left) and presence (right) of objects on consecutive trials. Mouse and cell number are indicated to the left of each pair of columns; grid score (G.s.) and peak rate are shown below each rate map. Cells in the upper three rows of the left pair of columns are examples of grid cells that expressed an extra field in the vicinity of at least one of the objects. Such effects were observed in 10 out of 40 grid cells. Only one grid cell passed the criteria for object-vector cells. Grid fields from the no-object trial mostly retained their firing locations when the object was added but in a few cases, single fields were moderately displaced. **b)** Colour-coded rate maps and head-direction tuning curves for 6 representative head-direction cells recorded with no object (left) and with an object (right) on consecutive trials. Mouse and cell number are indicated to the left; peak rate is shown below each rate map and head direction score (HD) and peak rate are shown below each head-direction tuning curve. A total of 38 out of 98 object-vector cells passed criteria for head-direction cells on no-object trials. Sharply tuned head-direction cells mostly failed to develop vector fields in the presence of objects (exemplified by the cells in the upper three rows). One of the few object-vector cells that also had sharp head-direction tuning is cell 398 in the fourth row. The majority of object-vector cells that passed as head-direction cells had only moderate head-direction tuning, and this tuning was clearly reduced when the object was introduced (exemplified by cells 351 and 266 in the two bottom rows; see also Fig. 5c,d).

